# Inner hair cell dysfunction in *Klhl18* mutant mice leads to low frequency progressive hearing loss

**DOI:** 10.1101/2021.03.09.434536

**Authors:** Neil J Ingham, Navid Banafshe, Clarisse Panganiban, Julia L Crunden, Jing Chen, Karen P Steel

**Affiliations:** Wolfson Centre for Age-Related Diseases, King’s College London, Guys Campus, London, SE1 1UL, UK

## Abstract

Age-related hearing loss in humans (presbycusis) typically involves impairment of high frequency sensitivity before becoming progressively more severe at lower frequencies. Pathologies initially affecting lower frequency regions of hearing are less common. Here we describe a progressive, predominantly low-frequency hearing impairment in two mutant mouse lines, carrying different mutant alleles of the *Klhl18* gene: a spontaneous missense mutation (*Klhl18^lowf^*) and a targeted mutation (*Klhl18^tm1a(KOMP)Wtsi^*). Both males and females were studied, and the two mutant lines showed similar phenotypes. Auditory brainstem response (ABR) thresholds (a measure of auditory nerve and brainstem neural activity) were normal at 3 weeks old but showed progressive increases from 4 weeks onwards. In contrast, distortion product otoacoustic emission (DPOAE) sensitivity and amplitudes (a reflection of cochlear outer hair cell function) remained normal in mutants. Electrophysiological recordings from the round window of *Klhl18^lowf^* mutants at 6 weeks old revealed 1) raised compound action potential thresholds that were similar to ABR thresholds, 2) cochlear microphonic potentials that were normal compared with wildtype and heterozygous control mice and 3) summating potentials that were reduced in amplitude compared to control mice. Scanning electron microscopy showed that *Klhl18^lowf^* mutant mice had abnormally tapering inner hair cell stereocilia in the apical half of the cochlea while their synapses appeared normal. These results suggest that Klhl18 is necessary to maintain inner hair cell stereocilia and normal inner hair cell function at low frequencies. *Klhl18* mutant mice exhibit an uncommon low frequency hearing impairment with physiological features consistent with Auditory Neuropathy Spectrum Disorder (ANSD).

**SIGNIFICANCE STATEMENT:** We describe a novel progressive hearing loss in *Klhl18* mutant mice that affects the lower frequencies of its’ hearing range. Investigation of two mutant alleles of this gene revealed primary inner hair cell defects affecting the neural output of the cochlea while outer hair cell function appeared normal. The tallest stereocilia of inner hair cells showed an abnormal tapering shape, especially notable in the apical half of the cochlear duct corresponding to the low frequency hearing loss. Our finding of a primary inner hair cell defect associated with raised thresholds for auditory brainstem responses combined with normal outer hair cell function suggests that Klhl18 deficiency and inner hair cell pathology may contribute to Auditory Neuropathy Spectrum Disorder in humans.

## INTRODUCTION

Progressive hearing loss with age is the most common sensory deficit in the human population and it can begin at any age. Genes found to be involved in hearing loss can give valuable insights into the molecular pathways and pathological processes leading to progressive deafness. Mouse mutants not only can provide valuable candidate genes for human deafness but also can reveal novel pathological mechanisms underlying hearing loss. We have used the mouse to identify multiple genes involved in hearing loss but it is clear that many more genes, and pathological mechanisms, remain to be discovered before we have a full understanding of the disease (Ingham et al., 2019).

In this report we explore the role of the *Klhl18* (Kelch-like family member 18) gene in hearing loss using two different mutations in the mouse. The first allele (termed *Klhl18^lowf^*) arose as a spontaneous missense mutation (V55F) predicted to have a damaging effect on protein structure (Lewis et al., 2021). These mutants have normal middle ears and no malformations of the inner ear were found (Lewis et al., 2021). The second allele is a targeted mutation, *Klhl18^tm1a(KOMP)Wtsi^*, exhibiting low frequency hearing loss in adult mice (Bowl et al., 2017, Ingham et al., 2019). Mice carrying the targeted allele have been subject to a broad-spectrum phenotyping screen as described by White et al. (2013). The only other significant phenotype found was a decreased volume and thickness of femur trabecular bone in female homozygous mutants (Ingham et al., 2019). Neither allele produced overt vestibular phenotypes and homozygotes are viable and fertile (Ingham et al., 2019; Lewis et al., 2021).

Kelch-like family member 18 is part of a 42-member superfamily of genes (Dhanoa et al., 2013). In the mouse, Klhl18 is a 574 amino acid protein containing a Bric-a-Brac, Tramtrack and Broad complex (BTB) domain, a BACK domain, and six Kelch β-propeller domain repeats which typically have roles in extracellular communication, cell morphology, and actin binding (Dhanoa et al., 2013). BTB domains are involved in protein-protein binding (Albagli et al., 1995; Perez-Torrado et al., 2006) and have been associated with a variety of cellular mechanisms, including cytoskeletal organization (Kang et al., 2004), voltage-gated potassium channel opening (Minor et al., 2000), transcriptional regulation (Melnick et al., 2000) and targeting of proteins for ubiquitination (Furukawa et al., 2003; Xu et al., 2003). *Klhl18* encodes an adaptor protein for the Cul3 ubiquitin ligase, providing specific targeting of Aurora-A for ubiquitination and subsequent initiation of mitotic entry (Moghe et al., 2012). Recently, Chaya et al. (2019) have implicated *Klhl18* in retinal photoreceptor function in mice through targeted ubiquitination of Unc119.

Here, we describe progressive elevation of auditory brainstem response (ABR) thresholds in *Klhl18* mutant mice from 4 weeks old. In contrast, *in vivo* measurements of distortion product otoacoustic emissions (DPOAEs) and cochlear microphonics (CM) indicated normal outer hair cell (OHC) function is maintained. The number of synaptic contacts between inner hair cells (IHCs) and afferent cochlear neurons was normal. However, IHC stereocilia displayed abnormal lengthening and tapering, especially affecting the apical half of the cochlear duct. These observations indicated the IHC as the primary site of the pathology. Summating potentials (SP), a sustained dc shift in voltage seen during sound exposure and thought to arise mostly from depolarisation of IHCs, were abnormally small in mutants supporting this suggestion.

## MATERIALS AND METHODS

### Ethics statement

Mouse studies were carried out in accordance with UK Home Office regulations and the UK Animals (Scientific Procedures) Act of 1986 (ASPA) under UK Home Office licences, and the study was approved by the King’s College London and Wellcome Trust Sanger Institute Ethical Review Committees. Mice were culled using methods approved under these licences to minimize any possibility of suffering.

### Mice

The mouse lines carrying the two alleles of *Klhl18* used in this study originated from the Wellcome Sanger Institute Mouse Genetics Project, both generated and maintained on a C57BL/6N genetic background. One was a spontaneous missense mutation, *Klhl18^lowf^* (Figure 1A), that occurred in a colony carrying a targeted mutation of *Mab21l4* (also known as *2310007B03Rik*). The mutation was detected through observation of occasional mice in this colony with raised ABR thresholds to low frequency stimuli that did not segregate with the targeted mutation in *Mab21l4.* The mutation was identified as a point mutation in *Klhl18* by positional cloning (g.9:110455454C>A), predicted to cause a Val55Phe amino acid change in the BTB domain of the protein (Lewis et al., 2020). The second was a targeted mutant allele (*Klhl18^tm1a(KOMP)Wtsi^*, referred to here as *Klhl18^tm1a^*, Figure 1B) carrying a promoter-driven knockout-first allele, with a large cassette inserted between exons 5 and 6, which interferes with transcription leading to knockdown of expression (Skarnes et al., 2011; White et al., 2013). The inserted cassette contains a β-galactosidase/LacZ reporter gene. Further details can be found at www.mousephenotype.org. Both *Klhl18* mutant lines produce homozygous offspring that are viable and fertile, and hearing impairment was inherited in a recessive manner. Both males and females were included as no difference in auditory function was noted between them. These mutant mice are available through the European Mouse Mutant Archive (EMMA).

**Figure 1.**
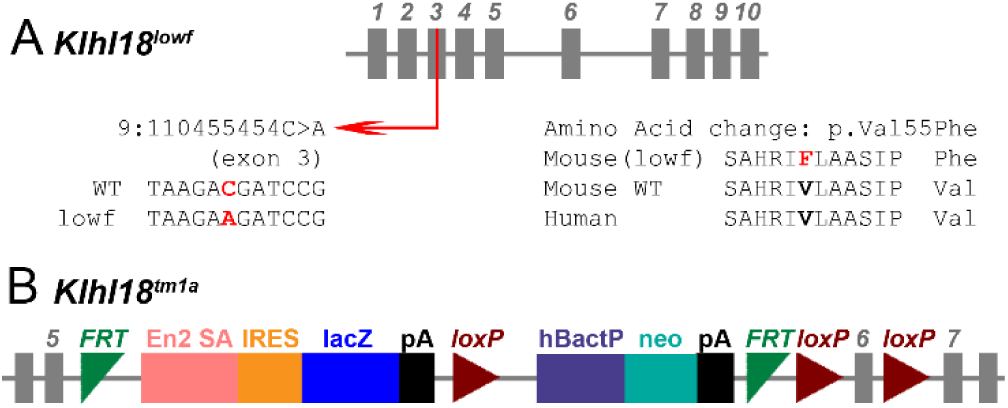
Schematic representations of the *Klhl18^lowf^* and *Klhl18^tm1a^* alleles. **A.** *Klhl18^lowf^* is formed by a single base pair change on Chr9, at position 110455454, in exon 3 of the gene, with the wildtype C replaced with the mutant A. This produces an amino acid substitution in the resultant protein with the conserved wildtype valine being replaced by phenylalanine at position 55 (see Lewis et al 2020 for further information). **B.** *Klhl18^tm1a^* was generated at the Wellcome Sanger Institute by insertion of a large DNA cassette before the critical exon (exon 6) of the gene, which interrupts transcription of the gene into mRNA. The cassette is composed of a number of components; En2 SA (engrailed2 splice acceptor), IRES (internal ribosome entry site), lacZ (beta-galactosidase reporter gene), pA (polyadenylation site), hBactP (beta-actin promotor part) and neo (neomycin resistance gene), which are flanked by FRT (Flp-FRT) and loxP recombination sites to produce the promoter driven Knockout First, reporter-tagged insertion with conditional potential allele (tm1a) of *Klhl18* (see Skarnes et al 2011 for further information).

### Genotyping

*Klhl18^lowf^* mice were genotyped by PCR amplification followed by restriction enzyme digest of the PCR product. The *Klhl18^lowf^* mutation is a C>A missense mutation in exon 10 of the gene (Lewis et al., 2021). We amplified genomic DNA in this region (forward primer sequence: GCACAATGGTAGGGGTTCAG; reverse primer sequence, GCAGTGTCGCTCAATATTTGTCTTTGTATTCTCTTTGGCCCACAGATTGGGGACCACAAGTTCAGTGCTCACCA G). The PCR primers used together with the point mutation generated a Bgl II restriction site (AGATCT) in the mutant PCR product while the corresponding wildtype allele sequence (AGATCC) was not recognised. BgI II was used to digest the PCR product yielding 2 sequences of 78 bp and 127 bp from the mutant allele and one sequence of 205 bp from the wildtype allele, and these products were identified by gel electrophoresis. In some cases, genotypes were obtained by sequencing of the PCR product amplified from the reverse primer. *Klhl18^tm1a^* mice were genotyped using a common forward primer (CCTGTGACAAGCAGTCTGAAGG), a wildtype reverse primer (TGCTAGGGAGTGAATCTAGGGC) and a mutant-specific reverse primer (CasR1 TCGTGGTATCGTTATGCGCC). Resulting band sizes were 524 bp for the wildtype product and 384 bp for the mutant product. Primers specific for the Neomycin resistance gene in the introduced cassette were also used to detect the presence or absence of the inserted DNA of the mutant allele (Figure 1; Skarnes et al., 2011; White et al., 2013).

### Anaesthesia

In experiments where mice were tested longitudinally at different ages, mice were anaesthetised by intra-peritoneal injection of 100 mg/kg Ketamine (Ketaset, Fort Dodge Animal Health) and 10 mg/kg Xylazine (Rompun, Bayer Animal Health) and recovery was promoted using 1 mg/kg atipamezole (Antisedan, Pfizer). For terminal experiments, mice were anaesthetised with intra- peritoneal urethane (0.1 ml / 10g bodyweight of a 20% w/v solution of urethane in water).

### Auditory Brainstem Response (ABR) recordings

Brainstem auditory evoked potentials were measured using the method described in detail by Ingham et al. (2011) and Ingham (2019). Anaesthetised mice were placed in a sound attenuating chamber (IAC Acoustics Limited) and subcutaneous recording needle electrodes (NeuroDart; Unimed Electrode Supplies Ltd, UK) were inserted on the vertex and overlying the left and right bullae. Responses were recorded to free-field calibrated broadband click stimuli (10 µs duration) and tone pips (5 ms duration, 1 ms onset and offset ramp) at frequencies between 6 and 42 kHz, at levels ranging from 0-95 dB SPL (in 5 dB steps) at a rate of 42.6 stimuli per second. Stimuli were generated via custom software on a RZ6 multifunction processor (Tucker Davis Technologies, TDT) and presented via a FF1 loudspeaker (TDT). Evoked responses were amplified, digitized, and bandpass filtered between 300-3000 Hz, using custom software and TDT hardware (RZ6 processor, RA4LI low impedance headstage, RA4PA preamplifier). ABR thresholds were defined as the lowest stimulus level to evoke a visually-detected waveform. ABR waveforms were analysed offline and the latency and amplitude of ABR waves 1-4 were plotted as a function of sound level.

### Distortion Product Otoacoustic Emissions (DPOAE) recordings

We made measurements of the 2f1- f2 DPOAE component in mice across a range of ages, either as a terminal experiment in different cohorts of mice, or as part of longitudinal experiments in the same animals at increasing ages. In all cases, measurements were made inside a sound attenuating chamber (IAC Ltd) with the mouse positioned on a heating blanket. In terminal experiments, urethane-anaesthetised mice had their left pinna and cartilaginous ear canal removed before a hollow conical speculum was positioned to give an unobstructed view of tympanic membrane. The DPOAE measurement probe (see later), with a small rubber gasket close to its’ tip was then positioned within the speculum. For longitudinal experiments, ketamine/xylazine-anaesthetised mice were placed in a prone position and the head was tilted approximately 45° such that the left ear was uppermost. The DPOAE probe assembly was positioned vertically, such that the probe tip was sitting just behind the tragus, with the probe pointing down towards the opening of the ear canal. The DPOAE probe assembly used here was comprised of a pair of EC1 electrostatic drivers (TDT) coupled to the guide tubes of an ER10B+ low noise DPOAE system (Etymotic Inc) via 5 cm plastic tubes.

Stimuli were generated and DPOAE responses recorded using a RZ6 multifunction processor (TDT), under the control of TDT BioSigRZ software (TDT). Continuous f1 and f2 tone stimuli were generated and presented via different EC1 drivers within the DPOAE probe. Frequencies for f2 were set to match the ABR tone-pip frequencies used (ie. 6, 12, 18, 24 and 30 kHz). The f2 tone was presented at a frequency 1.2x that of f1, and a level 10 dB SPL lower than f1. Sound pressure levels of the f2 stimulus ranged from -10 dB to 65 dB in 5 dB steps. ER10B+ microphone signals recording during stimulus presentation were digitised at a sampling rate of 195312.5 Hz on the RZ6 processor for online Fast Fourier Transformation (FFT) to yield a power spectrum containing the f1 and f2 stimulus components and the main 2f1-f2 DPOAE component of interest in this study.

From each FFT trace recorded, a number of parameters were calculated; the 2f1-f2 DPOAE amplitude, the mean noise-floor amplitude (as the average of the 20 spectral lines surrounding the DPOAE frequency) and two-times the standard deviation of the noise-floor mean. These values are plotted across stimulus level to produce plots from which the threshold of the DPOAE was defined as the lowest stimulus level when the DPOAE amplitude exceeded the 2 SDs above the recording noise-floor. From these data, we calculated mean DPOAE growth functions for each f2 stimulus frequency for control and mutant mice and mean DPOAE thresholds for these frequencies in control and mutant mice.

### Endocochlear Potential (EP) recordings

The positive potential within the scala media of *Klhl18^lowf^* mice was measured in urethane-anaesthetised mice as described previously (Steel and Barkway, 1989; Ingham et al., 2016). A reference electrode (Ag-AgCl pellet) was positioned under the skin of the neck. A small hole was made in the basal turn lateral wall and the tip of a 150 mM KCl-filled glass micropipette was inserted into the scala media. The EP was recorded as the differential potential between the tip of the glass electrode and the reference electrode.

### Round Window Response (RWR) recordings

Measurements of evoked potentials detected at the round window of the cochlea were made in *Klhl18^lowf^* mice aged 6 weeks. The methods used here were modified from those of Harvey and Steel (1992). In urethane-anaesthetised mice, following insertion of a tracheal cannula and placement in a custom-build head-holder, a subdermal needle reference electrode was inserted on the midline of the neck. A further subdermal needle electrode was inserted in the skin at the base of the tail to serve as a ground electrode. After removing the pinna, a small opening was made in the bulla and the exposed tip of a length of fine Teflon-coated silver wire was positioned on the round window membrane.

Auditory stimuli were generated using custom software and RZ6 multifunction processor (TDT) and presented via a MF1 magnetic loudspeaker (TDT) positioned 15 cm away from and directly opposite the exposed ear canal. Stimuli used were 20 ms duration tone pips, with a 1 ms ramp at the onset and offset, of the same frequencies used in the previous *in vivo* measurements (6, 12, 18, 24 and 30 kHz), with a fixed starting phase of 90°. These were presented from 0-95 dB SPL, at 10.65 stimuli / second, with the onset of the tone delayed by 5 ms after the onset of recording. For each sound level, 256 presentations of the tone pip contributed to the generation of an averaged evoked RWR. Signals detected at the Round Window were amplified (x1000 gain) and filtered (1 Hz high pass, 50 kHz low pass) using a DP311 differential amplifier (Warner Instruments). The amplified filtered signal was sampled at 97656.25 Hz by the RZ6 processor (TDT) and under software control, 80 ms snippets of the signal were used to form an averaged RWR to each combination of tone frequency and sound level.

### Analysis of Electrophysiological Responses from the Round Window of *Klhl18* mice

The RWRs recorded were analysed offline (Figure 6-1 extended data) using custom-written scripts in Matlab (v2019a, The Mathworks Inc., Natick MA, USA). Each RWR (Figure 6-1A extended data) was filtered into narrow bands to extract different components of the signal. A low pass filter (corner frequency, Fc, 3000Hz) was used to extract a combined CAP and Summating Potential (SP) response (Figure 6-1Bi extended data). The SP was measured as the mean amplitude of the waveform calculated over the 19-24 ms time window (equivalent to the 14-19 ms section of the tone burst stimulus). The SP could be either positive or negative in sign, relative to the zero baseline of the response (Figure 6-1Bii extended data). The amplitude of SP measured across stimulus level (from 0-95 dB) was plotted to form an input-output function (Figure 6-1Biii extended data). A bandpass filter (high pass Fc, 300 Hz; low pass Fc, 3 kHz) was used to extract the cochlear nerve Compound Action Potential (CAP) (Figure 6-1Ci extended data). The CAP was formed of significant negative and positive peaks to its waveform (labelled N and P, respectively in Figure 6-1Ci, 6-1Cii extended data). For each stimulus level, the N-P peak-to-peak amplitude was calculated and along with the N and P latencies following the onset of the tone pip were plotted to form input-output functions for these parameters (Figure 6-1Ciii, 6-1Civ extended data). A narrow bandpass filter centred at stimulus frequency (with low and high pass Fc’s of ± 100 Hz) was used to isolate the Cochlear Microphonic (CM) (Figure 6-1Di extended data). This filtered response was trimmed to a time window equivalent to the 2-18 ms section of the tone burst stimulus, and this steady-state section of the CM response was subject to Fast Fourier Transformation. The resultant power spectrum (Figure 6-1Dii extended data) was used to determine the amplitude of the CM component. The amplitude of the CM was converted to a dB scale, using 1 µV as a reference amplitude. The dB (re 1 µV) CM amplitude measured across stimulus level was plotted to form an input-output function of CM growth (Figure 6-1Diii extended data).

### Innervation and synaptic labelling with confocal microscopy imaging

Inner ears from 6 week old mice were fixed in 4% paraformaldehyde for 2 hours and decalcified in 0.1 M EDTA overnight at room temperature. Following fine dissection in PBS, the organ of Corti was permeabilised in 5% Tween20 in PBS for 40-60 minutes and incubated in blocking solution (0.45% Triton X-100, 10% normal horse serum in PBS) for 2 hours. Following blocking, the samples were incubated overnight at room temperature with shaking using primary antibodies in 0.36% Triton, 6% normal horse serum in PBS. For synaptic labelling, primary antibodies were mouse anti-GluR2 (1:200, MAB397, Emd Millipore) and rabbit anti-Ribeye (1:500, 192 103, Synaptic Systems). For neuron labelling, primary antibodies were mouse anti-CtBP2 (1:400, BD Transduction Laboratories 612044) to label pre-synaptic ribbons and IHC nuclei and chicken anti-Neurofilament-Heavy (1:800, Abcam ab4680) to label unmyelinated neural dendrites. Samples were then washed with PBS being incubated with secondary antibodies for 45-60 minutes in the dark. For synapses, secondary antibodies were either donkey anti-mouse IgG Alexa Fluor594 (A21203, ThermoFischer Scientific) or goat anti-mouse IgG2a Alexa Fluor488 (A21131, ThermoFischer Scientific) and goat anti-rabbit IgG Alexa Fluor546 (A11035, ThermoFischer Scientific). For neuron labelling, secondary antibodies were donkey anti-mouse Alexa Fluor594 (1:500, A-21203, Molecular Probes) and goat anti-chicken Alexa Fluor488 (1:300, A11039, Life technologies). Samples were then washed in PBS before mounting in ProLong Gold mounting medium with DAPI and stored at 4°C in the dark. Specimens were viewed using a Zeiss Imager 710 confocal microscope interfaced with ZEN 2010 software with a x63 objective and numerical aperture of 1.4. The whole length of the organ of Corti was examined and images collected at best-frequency regions based upon the frequency-place map of Müller et al. (2005). Z-stacks were taken at 0.25 µm intervals and maximum intensity projection images were generated. Brightness and contrast were normalised for the dynamic range in all images. The number of ribbon synapses per IHC was quantified by counting manually the co-localised Ribeye and GluR2 puncta in the confocal maximum projection images and dividing it by the number of DAPI-labelled IHC nuclei. An average of six IHCs per image were considered for the ribbon synapses counting using the cell-counter plugin in Fiji software.

### Scanning Electron Microscopy

Cochlear samples from *Klhl18^lowf^* mice aged P21 and P28 were fixed in 2.5% glutaraldehyde in 0.1 M sodium cacodylate buffer with 3 mM calcium chloride, dissected to expose the organ of Corti, then processed by a standard osmium tetroxide-thiocarbohydrazide (OTOTO) protocol (Hunter-Duvar, 1978). After dehydration, samples were subjected to critical point drying, mounted and viewed using a Jeol JSM-7800F Prime Schottky field emission scanning electron microscope. An overview of the cochlea was imaged to allow calculation of percentage distances along the cochlear duct to superimpose the frequency-place map (Muller et al., 2005), allowing subsequent imaging of consistent locations across different specimens. Images were assessed by three independent viewers who were blinded to genotype.

### β-Galactosidase staining

We used the β-galactosidase reporter gene in the inserted cassette of *Klhl18^tm1a^* mutants to investigate the distribution of expression. Inner ears of post-natal day (P) 13 *Klhl18^tm1a^* heterozygote mice were fixed in 4% paraformaldehyde for 2 hours at room temperature (RT) with rotation, washed twice with PBS for at least 30 minutes and decalcified in 0.1 M EDTA with rotation at room temperature until the bone was sufficiently soft (usually 2 or 3 days). Samples were washed for 30 minutes with a detergent solution (2 mM MgCl_2_; 0.02% NP-40; 0.01% sodium deoxycholate in PBS, pH 7.3). X-gal (Promega, cat.no. V394A) was added 1:50 to pre-warmed staining solution (5 mM K_3_Fe(CN)_6_ and 5 mM K_4_Fe(CN)_6_ in detergent solution), then the inner ears were stained at 37°C in the dark overnight. Following X-Gal staining, the samples were washed with PBS, dehydrated and embedded in paraffin wax. The samples were sectioned at 8 µm, counterstained using Nuclear Fast Red (VWR, cat.no. 342094W) and mounted using Eukitt quick-hardening mounting medium (Sigma-Aldrich). Sections were imaged using a Zeiss Axioskop microscope connected to AxioCam camera and interfaced with Axiovision 3.0 software.

## EXPERIMENTAL DESIGN AND STATISTICAL ANALYSES

We used heterozygotes as controls in most of the electrophysiological experiments because we did not see any obvious difference between heterozygotes and wildtype littermates. Both males and females were used as we did not detect any difference between their thresholds and both males and females were combined into a single group for their age and genotype. As the auditory phenotype changes with age of the homozygous mutants, we use littermates as controls and carefully age-matched mice at different ages; 2 weeks (P14 +/- 0 days); 3 weeks (P21 +/- 0 days); 4 weeks (P28 +/- 1 day); 6 weeks (P42 +/- 2 days); 8 weeks (P56 +/- 4 days); 14 weeks (P98 +/- 4 days).

All statistical comparisons were made using the analyses routines within Graphpad Prism (v8.4.2). Auditory Brainstem Response, Distortion Product Otoacoustic Emissions and Round Window Response data were analysed using a Mixed-effect Model, with either Sidak’s or Tukey’s multiple comparisons tests. Endocochlear Potential values were compared using a Kruskal-Wallis One-Way ANOVA. Counts of synaptic components were compared using a two-tailed unpaired t-test, with Welch’s correction. All tests were performed with alpha = 0.05. Statistical significance was determined at the level of p < 0.05.

## RESULTS

### *Klhl18* mutant mice showed progressive increases in ABR thresholds with age, particularly for low frequency stimuli

In control mice, ABR thresholds improved from 2 weeks old, through 3 weeks old to 4 weeks old (Figure 2A-C), indicating maturation of auditory sensitivity. In 2 week old mutants, ABR thresholds were slightly but statistically significantly elevated, compared to age-matched littermate controls (Figure 2A; Table 1-1 extended data). At 3 weeks old, ABR thresholds of mutant and control mice are comparable (Figure 2B). From the age of 4 weeks (Figure 2C-F), ABR thresholds for all stimuli were significantly higher in mutants than in age-matched controls (See Table 1-1 extended data). As the mice aged beyond 4 weeks, control mice maintain good ABR thresholds but there was a progressive elevation of thresholds from 4 weeks old to 14 weeks in the *Klhl18^lowf^* mutant mice, most pronounced at 6-18kHz (Figure 2C-F). A similar pattern of age-related threshold changes is seen in *Klhl18^tm1a^* mutant mice (Figure 2-2 extended data). There was no significant effect of sex on ABR thresholds of 6 weeks old control or mutant mice of either *Klhl18^lowf^* or *Klhl18^tm1a^* lines (Figure 2-3A,B extended data, Table 1-1 extended data).

**Figure 2.**
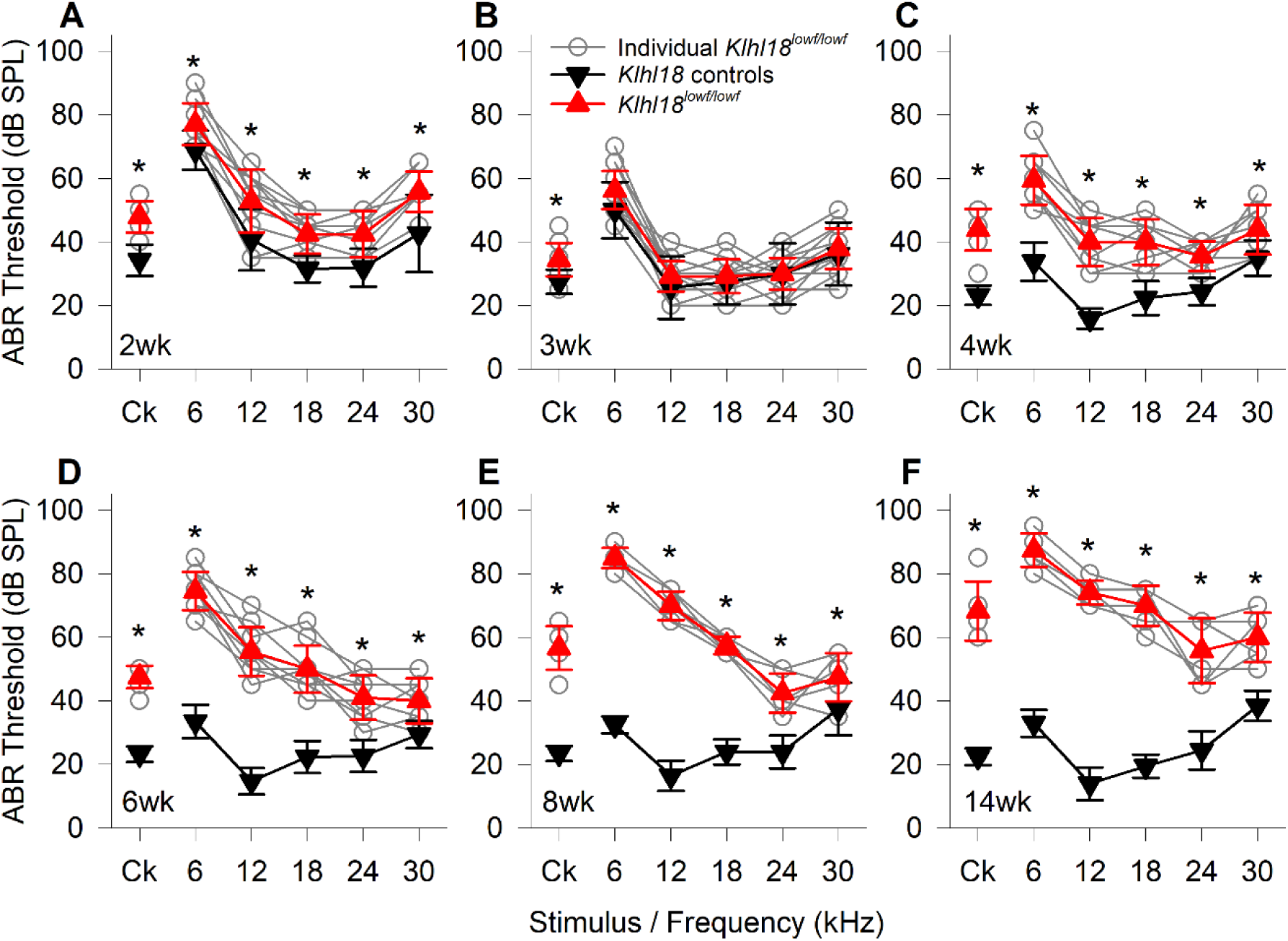
Age-Related changes in ABR thresholds of *Klhl18^lowf^* mice. A summary of ABR thresholds estimated from control (heterozygous) and mutant (homozygous) are shown for *Klhl18^lowf^* mice aged 2 weeks (**A**, n=13 control, n=12 mutant), 3 weeks (**B**, n=8 control, n=19 mutant), 4 weeks (**C**, n=17 control, n=9 mutant), 6 weeks (**D**, n=17 control, n=10 mutant), 8 weeks (**E**, n=10 control, n=6 mutant) and 14 weeks (**F**, n=10 control, n=6 mutant). Thresholds from individual mutant mice are indicated by grey circles. Mean threshold (±SD) are plotted for control mice as black down-triangles and for mutant mice as red up-triangles. Ck : Click stimulus. A mixed-effects model statistical analysis between control and mutant thresholds gave the following results; 2 weeks, F (1, 66) = 108.017, p=2.000x10^-15^; 3 weeks, F (1, 150) = 9.233, p=0.003; 4 weeks, F (1, 144) = 370.051, p=1.000 x10^-15^; 6 weeks, F (1, 150) = 976.621, p=1.000 x10^-15^; 8 weeks, F (1, 30) = 1268.834, p=1.000 x10^-15^; 14 weeks, F (1, 30) = 1464.112, p=1.000 x10^-15^. Sidak’s multiple comparisons test was used to examine the difference between control and mutant thresholds for each stimulus. Significant differences are given in Table 1-1 and indicated here by *. Equivalent data obtained from mice carrying the *Klhl18^tm1a^* allele are shown in figure 2-2 (extended data).

**Figure 3.**
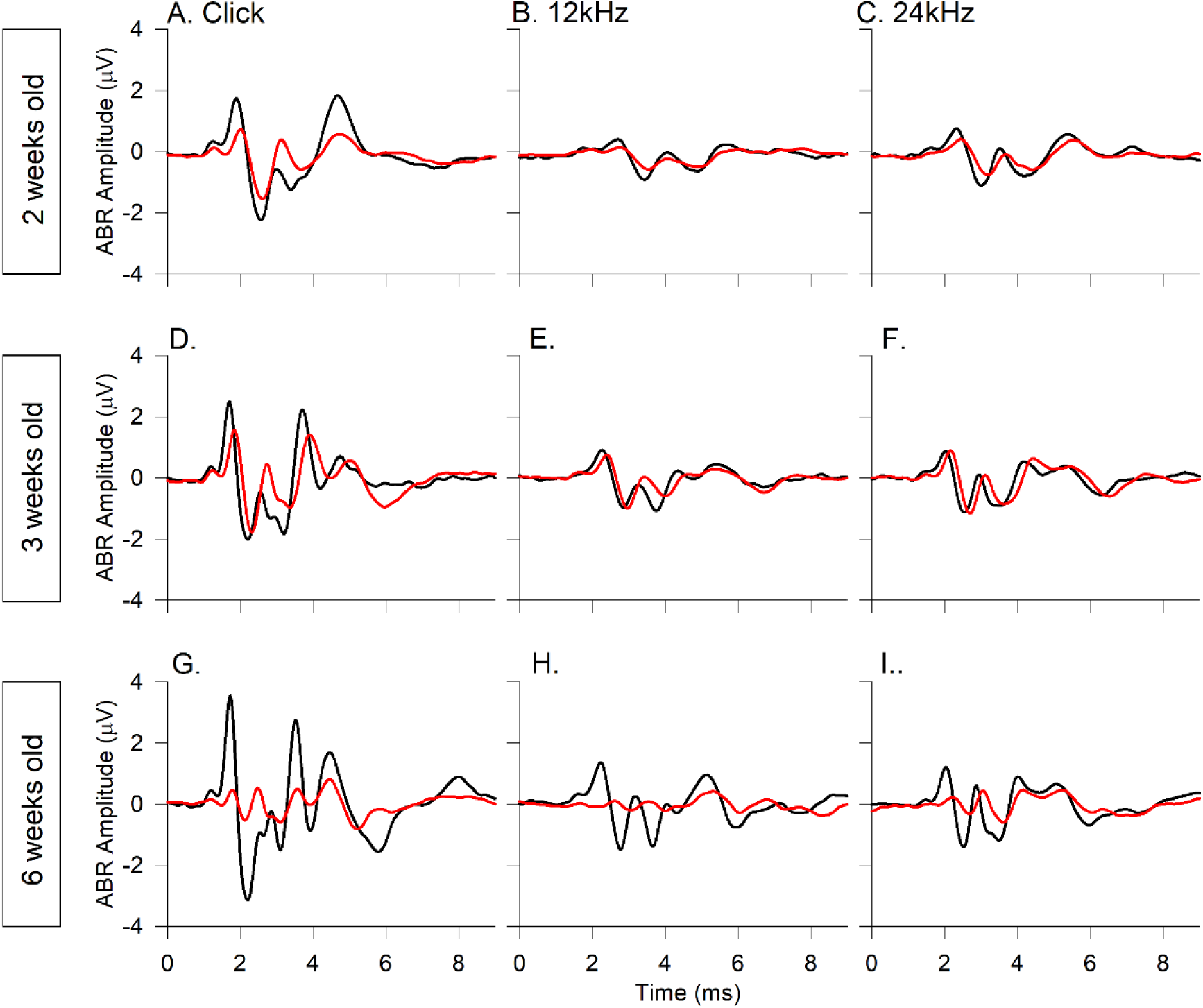
Mean ABR waveforms of *Klhl18^lowf^* mice at 65 dB SPL. Group averaged ABR waveforms, evoked by 65 dB SPL stimuli, are plotted for mice aged 2 weeks (**A-C**, n=13 control, n=12 mutant), 3 weeks (**D-F**, n=8 control, n=19 mutant), 6 weeks (**G-I**, n=17 control, n=10 mutant), in response to click stimuli (**A,D,G**), 12 kHz stimuli (**B,E,H**) and 24 kHz stimuli (**C,F,I**). Mean amplitude waveforms are plotted for the same control mice (black lines) and mutant mice (red lines) described in figure 2. Equivalent data obtained from mice carrying the *Klhl18^tm1a^* allele are shown in figure 3-2 (extended data).

### *Klhl18* mutant mice showed abnormal ABR waveform shapes

Group averaged ABR waveforms for selected age groups and frequency stimuli (all at 65 dB SPL) are plotted in Figure 3. For mice aged 2 weeks old, mean waveforms evoked by click stimuli, and 12kHz and 24 kHz tone pips (plotted in Figure 3 A-C), suggest that responses from *Klhl18* mutant mice had smaller amplitudes than controls. At 3 weeks old, when ABR threshold sensitivity was comparable across genotypes, the averaged waveforms for these stimuli were more comparable (Figure 3 D-F); the mean click evoked response was somewhat smaller in mutants, but the mean tone evoked response had a more comparable amplitude in both control and mutant mice. These mean waveform amplitudes from 2-3 weeks old mice were in contrast to those from older mice. At 6 weeks old, when mutant mice exhibited a clear loss of threshold sensitivity, the averaged waveforms of mutant mice evoked by click, 12 kHz and 24 kHz stimuli were much smaller than comparable averages taken from responses of control mice (Figure 3 G-I). Similar observations were obtained from *Klhl18^tm1a^* mutant mice (Figure 3-1 extended data). To account for the reduced auditory sensitivity in the *Klhl18* mutant mice, we also plotted mean waveform amplitudes evoked by stimuli at 20 dB sensation level (dB SL, ie dB above threshold). These group averages showed a similar pattern. ABR waveforms were similar in control and mutant mice aged 2-3 weeks, but were more abnormal in 6 weeks old mutants, where the averaged wave 1 amplitude was reduced in amplitude (Figures 3-2, 3-3 extended data).

### ABR Wave 1 showed reduced amplitude and longer latency in *Klhl18* mutant mice

ABR wave 1 amplitudes (measured as wave P1 – N1 peak to peak amplitude) and P1 latencies were plotted as a function of stimulus level (Figure 4. See also Figure 2-1, extended data). The mild threshold elevation in 2-week old *Klhl18^lowf^* mutants was reflected in a shift to the right of growth curves (Figure 4A-C, J- L). In mutant mice aged 3 weeks old, wave 1 amplitude was similar to controls (Figure 4D-F). However, P1 latency maintained a slight increase in mutants compared to age-matched controls (Figure 4M-O). By 6 weeks old, when ABR thresholds were clearly raised in mutants, much larger changes in ABR wave 1 amplitude and latency were noted (Figure 4G-I, P-R). Similar patterns of amplitude and latency changes were observed in *Klhl18^tm1a^* mutant mice (Figure 4-1 extended data). Amplitudes and latencies of ABR wave 1 in *Klhl18^tm1a^* heterozygotes were similar to their wildtype littermates (Figure 4-2 extended data).

**Figure 4.**
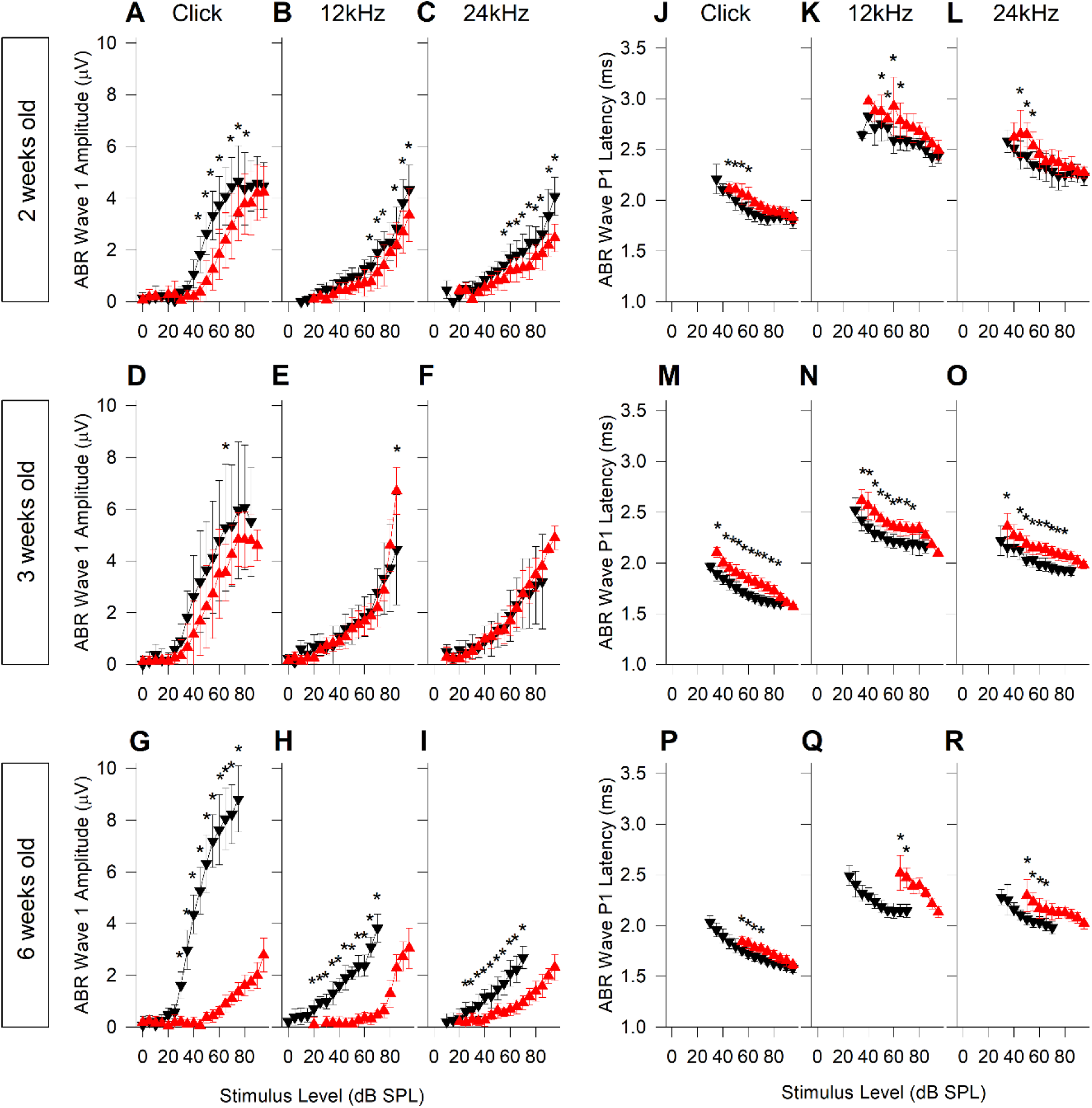
ABR wave 1 amplitude and latency input-output functions in *Klhl18^lowf^* mice. Mean ABR wave 1 P1-N1 peak-to-peak amplitude (±SD) as a function of stimulus level is plotted for mice aged 2, 3 and 6 weeks old are plotted in **A-C** (n=13 controls, n=12 mutants), **D-F** (n=8 controls, n=19 mutants) and **G-I** (n=17 controls, n=10 mutants), respectively. Results are plotted for click stimuli (**A,D,G**), 12 kHz tones (**B,E,H**) and 24 kHz tones (**C,F,I**). Data from control mice are plotted as black down-triangles. Data from mutant mice are plotted as red up-triangles. Mean ABR wave P1 latency (±SD) as a function of stimulus level is plotted for the same mice aged 2, 3 and 6 weeks old are plotted in **J-L**, **M-O** and **P- R**, respectively. Results are plotted for click stimuli (**J,M,P**), 12 kHz tones (**K,N,Q**) and 24 kHz tones (**L,O,R**). A mixed-effects model statistical analysis between control and mutant thresholds gave the following results; Wave 1 amplitude evoked by clicks at 2 weeks, F (1, 174) = 105.845, p=1.000 x10^-15^; 3 weeks, F (1, 117) = 50.042, p=1.186 x10^-10^; 6 weeks, F (1, 260) = 3056.781, p=1.000 x10^-15^; Wave 1 amplitude evoked by 12 kHz at 2 weeks, F (1, 124) = 77.410, p=1.000x10^-14^; 3 weeks, F (1, 108) = 1.147, p=0.287; 6 weeks, F (1, 71) = 731.550, p=1.000 x10^-15^; Wave 1 amplitude evoked by 24 kHz at 2 weeks, F (1, 126) = 129.325, p=1.000 x10^-15^; 3 weeks, F (1, 339) = 0.082, p=0.774; 6 weeks, F (1, 72) = 192.808, p=1.000 x10^-15^; P1 latency evoked by clicks at 2 weeks, F (1, 240) = 69.503, p=6.000 x10^-15^; 3 weeks, F (1, 68) = 241.820, p=1.000 x10^-15^; 6 weeks, F (1, 32) = 56.435, p=1.489 x10^-8^; P1 latency evoked by 12 kHz at 2 weeks, F (1, 190) = 66.516, p=4.600 x10^-14^; 3 weeks, F (1, 229) = 187.329, p=1.000 x10^-15^; 6 weeks, F (1, 7) = 55.045, p=1.470 x10^-4^; P1 latency evoked by 24 kHz at 2 weeks, F (1, 94) = 52.252, p=1.280 x10^-10^; 3 weeks, F (1, 232) = 150.525, p=1.000 x10^-15^; 6 weeks, F (1, 30) = 88.403, p=1.884 x10^10^. Sidak’s multiple comparisons test was used to examine the difference between control and mutant data for each stimulus. Significant differences are given in Table 1-1 and indicated here by *. Equivalent data obtained from mice carrying the *Klhl18^tm1a^* allele are shown in figure 4-1 (extended data).

### Normal DPOAE responses in *Klhl18* mutant mice

At 2 weeks old, there was a small statistically significant reduction in DPOAE amplitudes and elevation in DPOAE thresholds (for f2 frequencies of 12-30 kHz) in *Klhl18^lowf^* mutants, similar to the small differences in ABR responses recorded at this age (Figure 5A-F). In mice aged 3 weeks old, DPOAE responses were the same in *Klhl18^lowf^* mutants as in littermate controls, again mimicking the ABR thresholds at 3 weeks old (Figure 5G-L; no significant interaction of either DPOAE amplitude or threshold with genotype; See Table 1-1 extended data). However, in contrast to the significant elevation in ABR thresholds by the age of 6 weeks (Figure 3), there was no difference between DPOAE responses of control and mutant mice at this age Figure 5M-R; no significant interaction of either DPOAE amplitude or threshold with genotype; See Table 1-1 extended data). Furthermore, no differences were found in DPOAEs in *Klhl18^tm1a^* mutant mice up to 6 weeks old (extended data Figure 5-1). There was no significant effect of sex on DPOAE thresholds of 6 weeks old control or mutant mice of either *Klhl18^lowf^* or *Klhl18^tm1a^* lines (Figure 2-3C,D extended data, Table 1-1 extended data).

**Figure 5.**
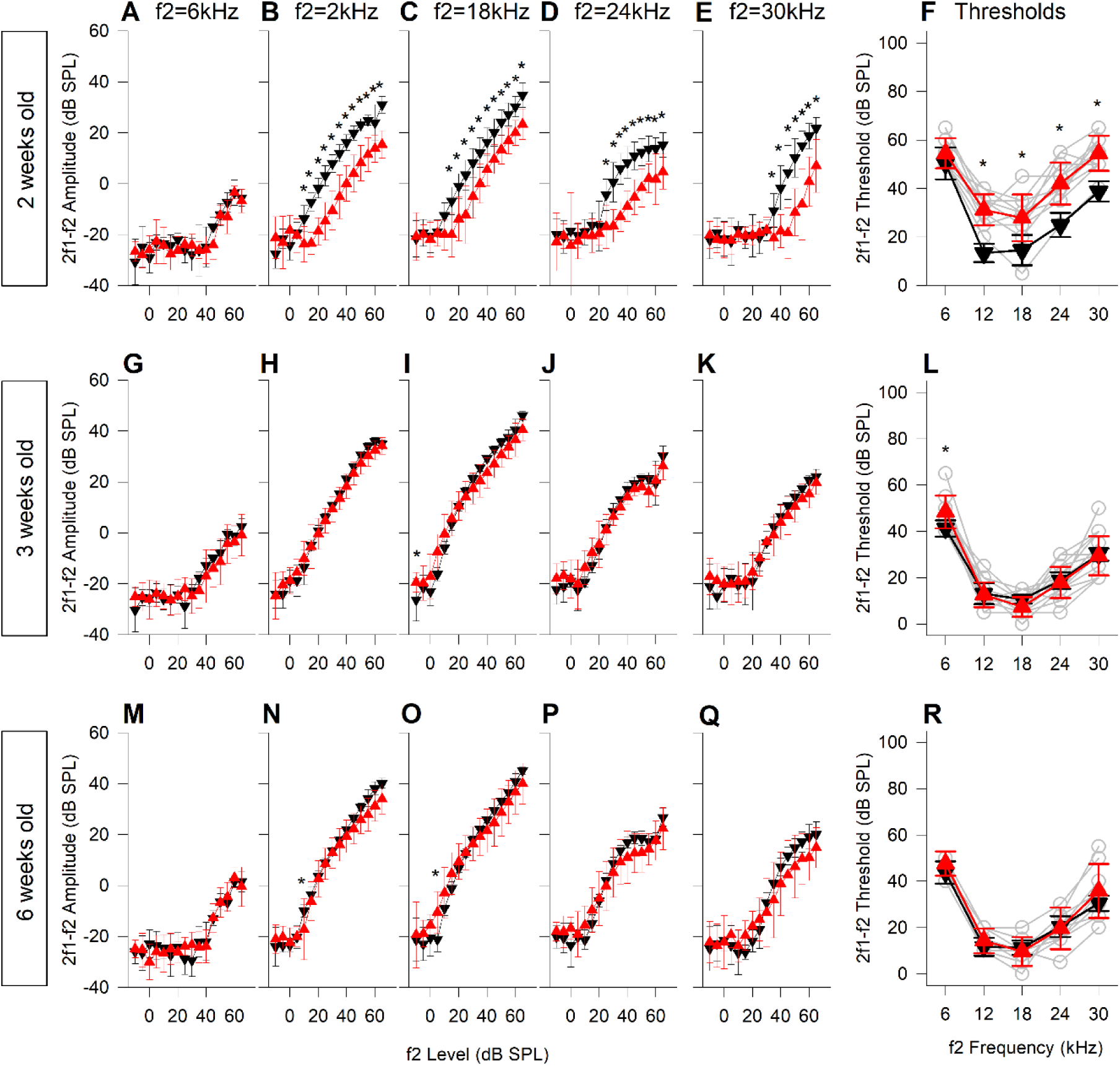
DPOAE growth and thresholds in *Klhl18^lowf^* mice. DPOAE results from mice aged 2, 3 and 6 weeks old are plotted in **A-F** (n=13 controls, n=12 mutants), **G-L** (n=8 controls, n=19 mutants) and **M- N** (n=9 controls, n=13 mutants), respectively. Data from control mice are plotted as black down- triangles. Data from mutant mice are plotted as red up-triangles. The mean (±SD) amplitude of the 2f1-f2 DPOAE (dB SPL) is plotted as a function of f2 stimulus level (dB SPL) for f2 frequencies of 6 kHz (**A,G,M**), 12 kHz (**B,H,N**), 18 kHz (**C,I,O**), 24 kHz (**D,J,P**) and 30 kHz (**E,K,Q**). Mean threshold (±SD) of the 2f1-f2 DPOAE (derived from individual growth functions, eg shown in Figure 5-1 extended data) and plotted in **F**, **L** and **R** for mice aged 2 weeks, 3 weeks and 6 weeks respectively. In addition to the mean data, thresholds from individual mutant mice are plotted as grey open circles. A mixed-effects model statistical analysis between control and mutant data gave the following results; DPOAE amplitude evoked by an f2 of 6 kHz at 2 weeks, F (1, 368) = 2.229, p=0.136; 3 weeks, F (1, 112) = 2.312, p=0.131; 6 weeks, F (1, 128) = 0.003, p=0.956; DPOAE amplitude evoked by an f2 of 12 kHz at 2 weeks, F (1, 176) = 375.214, p=1.000 x10^-15^; 3 weeks, F (1, 400) = 2.775, p=0.096; 6 weeks, F (1, 320) = 16.757, p=5.385 x10^-5^; DPOAE amplitude evoked by an f2 of 18 kHz at 2 weeks, F (1, 176) = 258.928, p=1.000 x10^-15^; 3 weeks, F (1, 112) = 1.548, p=0.216; 6 weeks, F (1, 320) = 0.167, p=0.683; DPOAE amplitude evoked by an f2 of 24 kHz at 2 weeks, F (1, 176) = 181.738, p=1.000 x10^-15^; 3 weeks, F (1, 112) = 0.066, p=0.798; 6 weeks, F (1, 320) = 1.050, p=0.306; DPOAE amplitude evoked by an f2 of 30 kHz at 2 weeks, F (1, 368) = 130.537, p=1.000 x10^-15^; 3 weeks, F (1, 112) = 0.543, p=0.462; 6 weeks, F (1, 320) = 3.551, p=0.060; DPOAE thresholds evoked across all stimulus frequencies at 2 weeks, F (1, 55) = 165.597, p=1.000 x10^-15^; 3 weeks, F (1, 35) = 0.810, p=0.374462; 6 weeks, F (1, 100) = 2.592, p=0.111; Sidak’s multiple comparisons test was used to examine the difference between control and mutant data for each stimulus. Significant differences are given in Table 1-1 and indicated here by *. Equivalent data obtained from mice carrying the *Klhl18^tm1a^* allele are shown in figure 5-2 (extended data).

**Figure 6.**
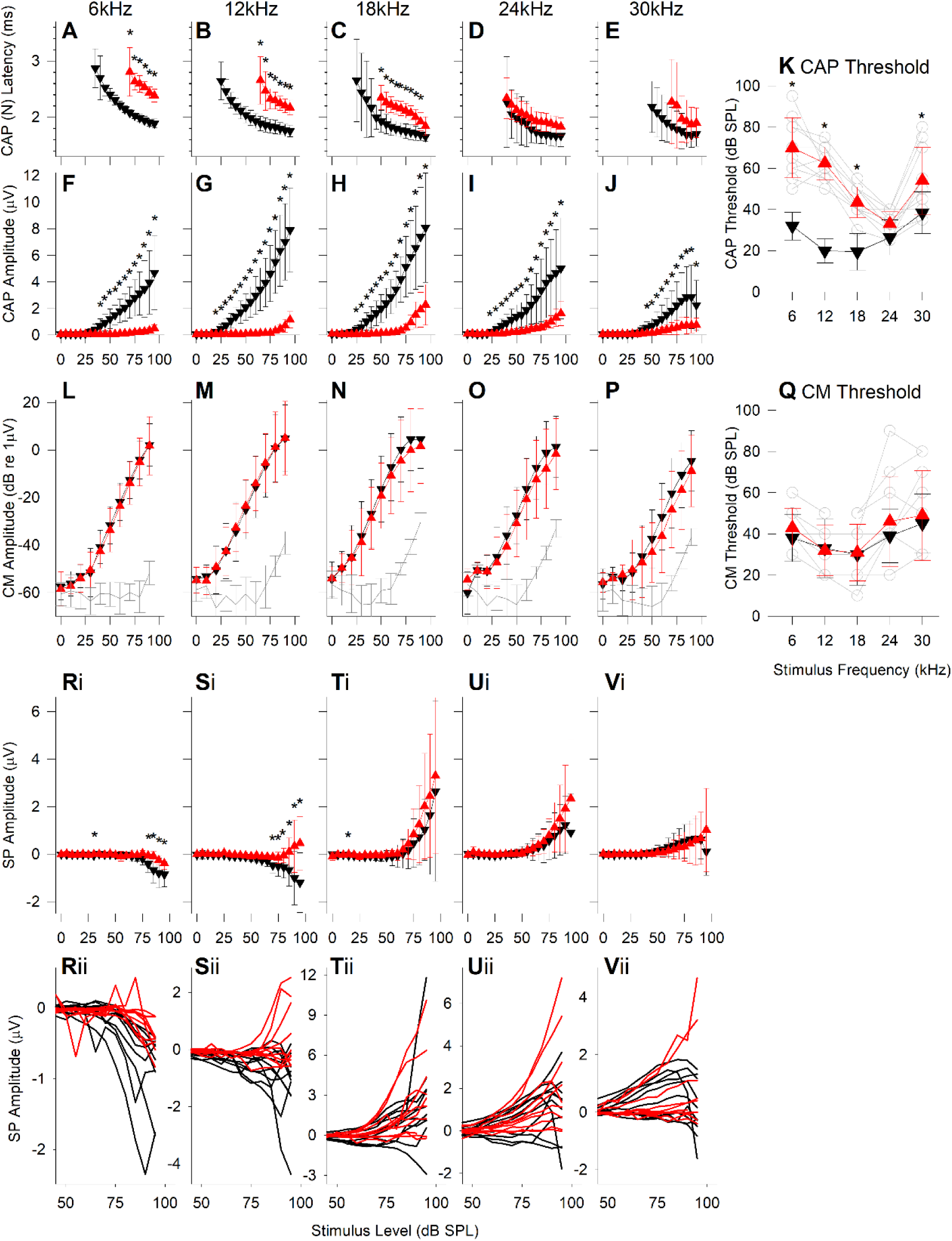
Round Window Response measurements from 6 weeks old *Klhl18^lowf^* mice. RWRs were obtained from 10 wildtype (*Klhl18^+/+^*, aged 41.8 ± 0.6 days), 7 heterozygote (*Klhl18^+/lowf^* aged 42.6 ± 1.1 days) and 10 homozygote mutant (*Klhl18^lowf/lowf^* aged 41.3 ± 1.2 days) mice. Compound Action Potential latency (of wave N) and amplitude (N-P amplitude) are plotted in **A-E** and **F-J**, respectively, for potentials measured in response to tones of 6, 12, 18, 24 and 30 kHz. Data are plotted as mean ± SD for wildtype control mice (black down-triangles) and homozygote mutant mice (red up-triangles). **K**. Mean ± SD of the CAP threshold is plotted against stimulus frequency for wildtype control mice (black down-triangles) and homozygote mutant mice (red up-triangles). Open circles and grey lines indicate CAP thresholds for individual mutant mice. **L-P** plot mean (± SD) Cochlear Microphonic amplitude (dB re 1 µV) against stimulus level (dB SPL) for wildtype mice (black down-triangles) and homozygote mutant mice (red up-triangles). We were unable to completely remove stimulus artefact from the RWRs recorded in our system. To account for this, we assessed the magnitude of the stimulus artefact across frequency and sound level, from analysis of measurements taken after the mouse was removed the system and the recording electrodes placed in 150 mM KCl. The biological CM was orders of magnitude greater in amplitude than the stimulus artefact and thus it was distinguishable from the artefact. The artefact estimated across all experiments, across stimulus frequency and level are plotted as grey lines (with SD error bars). **Q**. Mean ± SD of the estimated CM threshold is plotted against stimulus frequency for wildtype control mice (black down-triangles) and homozygote mutant mice (red up-triangles). Open circles and grey lines indicate CM thresholds for individual mutant mice. **Ri-Vi** plot mean (± SD) Summating Potential amplitude (µV) against stimulus level (dB SPL) for wildtype mice (black down-triangles) and homozygote mutant mice (red up-triangles). **Rii-Vii** plot SP IOFs for individual control (black lines) and mutant (red lines) mice. A mixed-effects model statistical analysis between wildtype, heterozygote (see extended data figure 6-4) and homozygote data gave the following results; CAP N1 latency evoked by 6 kHz, F (1.5, 102.6) = 195.805, p=1.000 x10^-15^; 12 kHz, F (1.8, 147.0) = 172.909, p=1.000 x10^-15^; 18 kHz, F (1.7, 205.4) = 157.252, p=1.000 x10^-15^; 24 kHz, F (1.3, 118.4) = 5.205, p=1.603 x10^-2^; 30 kHz, F (1.6, 111.4) = 10.272, p=3.020 x10^-4^; CAP Amplitude evoked by 6 kHz, F (1.4, 324.6) = 108.900, p=1.000 x10^-15^; 12 kHz, F (1.6, 232.8) = 217.219, p=1.000 x10^-15^; 18 kHz, F (1.8, 262.7) = 142.609, p=1.000 x10^-15^; 24 kHz, F (1.5, 223.7) = 65.786, p=1.000 x10^-15^; 30 kHz, F (1.4, 205.2) = 47.295, p=1.330 x10^-13^; CAP threshold across all stimuli, F (1.9, 70.7) = 136.094, p=1.000 x10^-15^; CM Amplitude evoked by 6 kHz, F (1.9, 144.9) = 0.672, p=0.509; 12 kHz, F (1.8, 130.8) = 0.712, p=0.475; 18 kHz, F (1.8, 16.6) = 0.354, p=0.690; 24 kHz, F (1.8, 134.6) = 3.155, p-0.051; 30 kHz, F (1.8, 132.0) = 3.425, p=4.905 x10^-2^; CM threshold across all stimuli, F (1.8, 67.4) = 0.976, p=0.374; SP Amplitude evoked by 6 kHz, F (1.9, 276.9) = 23.392, p=1.251 x10^-9^; 12 kHz, F (1.8, 272.8) = 32.512, p=1.366 x10^-12^; 18 kHz, F (1.7, 250.2) = 5.553, p=6.707 x10^-3^; 24 kHz, F (1.6, 236.7) = 4.295, p=2.169 x10^-2^; 30 kHz, F (1.5, 226.8) = 3.102, p=0.060. Sidak’s multiple comparisons test was used to examine the difference between control and mutant data for each stimulus. Significant differences are given in Table 1-1 and indicated here by *.

### Endocochlear Potential (EP) was normal in *Klhl18^lowf^* mutant mice

Endocochlear potential was measured in 6 weeks old *Klhl18^lowf^* mutant and littermate control mice. EP was 122.4 ± 5.4 mV (mean ± SD) in 6 wildtype mice (aged 42.0 ± 0 days), 121.2 ± 5.3 mV in 6 heterozygote mice (aged 42.3 ± 1.2 days) and 123.4 ± 5.6 mV in 7 homozygous mutant mice (aged 42.1 ± 0.9 days). There was no significant difference in EP between mice of these 3 genotypes (See Table 1-1 extended data).

### Round Window Responses (RWRs) match ABR and DPOAE responses

We analysed three different responses extracted from recordings from the round window in 6-week old *Klhl18^lowf^* mutants and littermate controls; the Compound Action Potential (CAP), the Cochlear Microphonic (CM) and the Summating Potential (SP) (Figure 6). Extraction of the different potentials is illustrated in extended data Figure 6-1. Examples of RWRs and their components are shown in extended data Figures 6-2 and 6-3 for a *Klhl18^+/+^* wildtype mouse and a *Klhl18^lowf/lowf^* mutant mouse, respectively, for 18kHz tone stimuli.

Compound action potentials (CAP), reflecting auditory nerve activity, had thresholds and amplitudes very similar to those seen for ABRs, with low frequencies more severely affected than high frequencies in mutants (compare Figure 6K with Figure 2D, and Figure 6G,I with Figure 4H,I). Latencies of CAP responses were longer in mutants than in controls (Figure 6A-E) but the shift in latency at each frequency matched the shift in threshold so the latencies appear to be normal when adjusted for hearing level. There was no significant effect of sex on CAP thresholds of wildtype or *Klhl18^lowf^* mutant mice (Figure 2-3E extended data, Table 1-1 extended data).

Cochlear microphonics (CM), reflecting mostly outer hair cell activity, showed no significant difference in thresholds or amplitudes in *Klhl18^lowf/lowf^* compared with *Klhl18^+/+^* mice (Figure 6L-Q, Table 1-1 extended data). This matched our findings from DPOAEs, which were also normal in the mutants. There was no significant effect of sex on CM thresholds of wildtype or *Klhl18^lowf^* mutant mice (Figure 2-3F extended data, Table 1-1 extended data).

Summating potentials (SP) are a dc shift in the round window recording sustained for the duration of a toneburst and are thought to reflect hair cell depolarisation, particularly of basal turn IHCs (Russell et al., 1986; Cody and Russell, 1987). They can be negative or positive in polarity, with negative SP usually detected at lower frequencies and higher stimulus intensities. We found these responses to be variable between individual mice, whatever their genotype (Figure 6Rii-Vii). Despite the variability, SP amplitudes were significantly different in mutants at 6 and 12 kHz, while there was no difference in SP at higher frequencies (See Figure 6 and Table 1-1 extended data). No significant differences between wildtypes (*Klhl18^+/+^*) and heterozygotes (*Klhl18^+/lowf^*) were found in any of the round window measures analysed (Figure 6-4 and Table 1-1 extended data).

### Normal innervation of the cochlea

The presence of abnormal ABRs and CAPs together with normal DPOAE and CM responses and normal EPs suggests that *Klhl18* mutant mice exhibit an IHC or neural hearing loss, so we looked in more detail at the neural elements of the organ of Corti. We used neurofilament immunolabelling in whole mount organ of Corti preparations to examine the dendrites of cochlear neurons in 6-week old *Klhl18^lowf^* mutants and littermate controls in regions corresponding to 6, 18 and 30 kHz characteristic frequencies (Figure 7). Confocal imaging showed a normal pattern of innervation in all regions examined. To examine synaptic organization in *Klhl18^lowf^* mice, we immunolabelled GluR2 (a post-synaptic glutamate receptor subunit) and Ribeye (a component of presynaptic ribbon synapses) in the organ of Corti of 6 week old mice (Figure 8). We found no significant differences (see Table 1-1 extended data) in numbers of ribbons or postsynaptic densities or co-localised labelled puncta (indicating synapses) in these mutants compared with littermate controls, either in regions exhibiting raised ABR thresholds (12 kHz region) or relatively normal ABR thresholds (24 kHz region).

**Figure 7.**
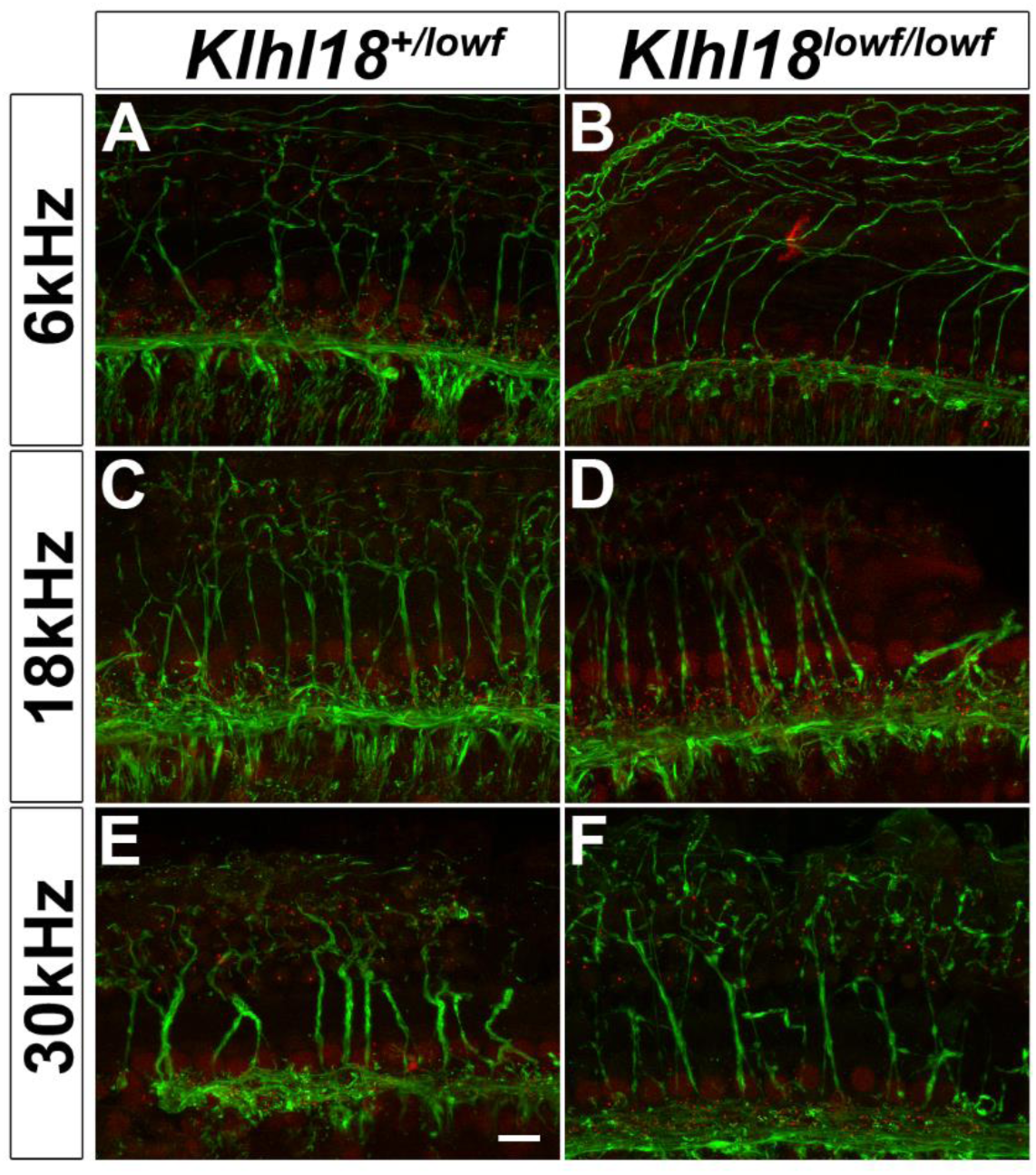
Auditory Nerve innervation of the organ of Corti at 6 weeks old. Representative confocal microscopy images of the organ of Corti at the 6 kHz (**A,B**), 18 kHz (**C,D**) and 30 kHz (**E,F**) regions of the cochlea from a sample of 7 ears from 5 *Klhl18^+/lowf^* mice (**A,C,E**), and 8 ears from 6 *Klhl18^lowf/lowf^* mice (**B,D,F**). CtBP2-positive regions (IHC cell nuclei and ribbon synapses) are immunostained in red; Neurofilament-positive regions (nerve fibres) are immunostained in green. The scale bar shown in (**E**) represents 10µm.

**Figure 8.**
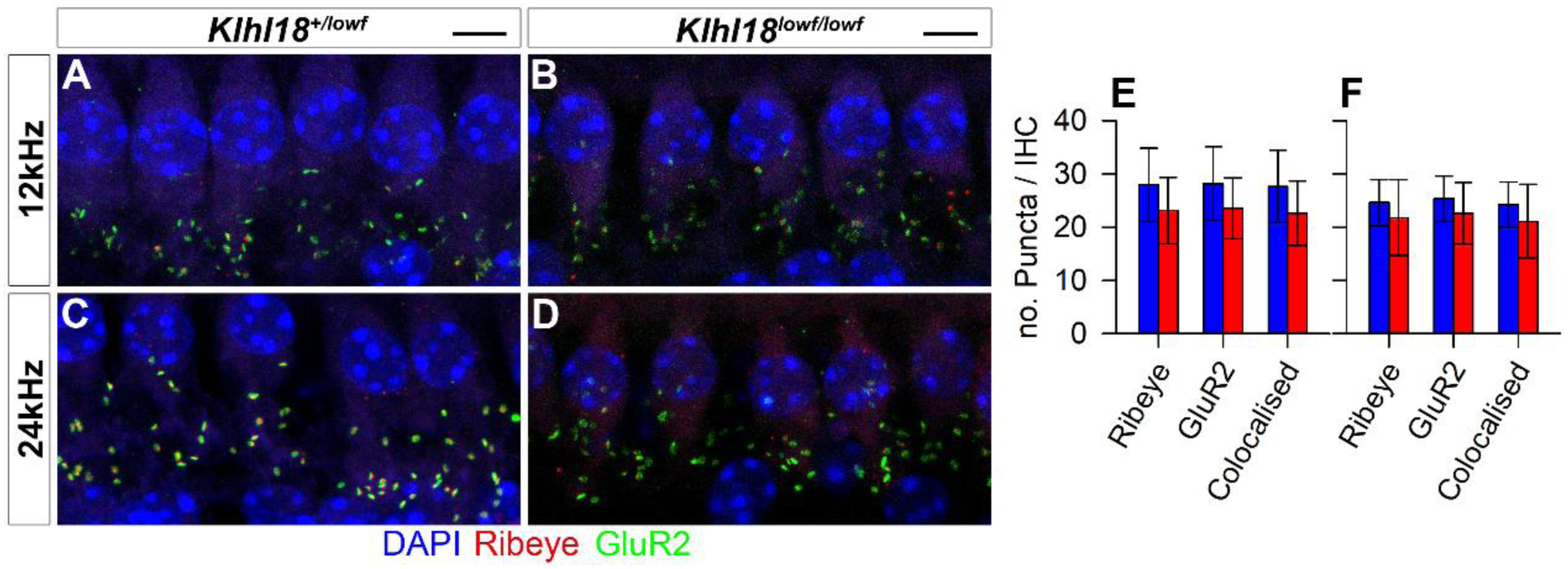
IHC synapses in Klhl18 mutant mice at 6 weeks old. Representative confocal microscopy images of Inner Hair Cells at the 12 kHz (**A,B**) and 24 kHz (**C,D**) regions of the cochlea from a sample of 7 *Klhl18^+/lowf^* mice (**A,C**), and 7 *Klhl18^lowf/lowf^* mice (**B,D**). A row of Inner Hair Cell nuclei can be seen across the upper portion of panels A-D, stained blue with DAPI. Below this row, clusters of immunostained puncta are visible. Green puncta show immuno-positive staining for the post-synaptic receptor subunit GluR2. Red puncta show immune-positive staining for Ribeye, a protein component of the ribbon synapse apparatus found in Inner Hair Cells. Yellow staining represents overlap between GluR2 and Ribeye (colocalised) staining as is presumed to represent a functional synapse between the Inner Hair Cells and the Auditory Nerve. Scale bars indicate 5µm. (**E,F**) Counts of Ribeye-positive, GluR2-positive and colocalised Ribeye and GluR2-positive puncta are plotted for the 12 kHz region (E) and the 24 kHz region (F). Data are plotted as bars representing the mean count per IHC (error bars indicate the standard deviation) for control (*Klhl18^+/lowf^*, blue bars) mice and homozygous mutant mice (*Klhl18^lowf/lowf^*, red bars).

### Tapering stereocilia in *Klhl18^lowf^* mutants

We examined the apical surface of the organ of Corti by scanning electron microscopy to look for any signs of hair cell degeneration or other defects. Homozygotes, heterozygotes and wildtypes were studied at two ages, P21 and P28, corresponding to stages when ABR thresholds were normal (P21) and when thresholds had started to increase in homozygotes (P28). In general, the organization of the sensory epithelium, the general patterning of inner and outer hair cells within the epithelium, and the appearance of all the associated supporting cell types were normal in *Klhl18^lowf^* mutant mice. There was no evidence seen of hair cell degeneration in the mutants, although we did not analyse the lower basal turn.

However, we did observe stereocilia abnormalities in the mutants that were most pronounced in the apical turn. Surprisingly we found similar defects at both 3 and 4 weeks old. The most severe defects were found in inner hair cells which showed frequent tapering of the tallest stereocilia towards their tips and they also appeared to be longer and thinner than normal (Figure 9 A-H). This feature was variable between hair bundles and also between individual stereocilia within a single bundle. The shorter two rows of stereocilia had a more normal appearance. Outer hair cell stereocilia of mutants looked close to normal, but some bundles appeared slightly more irregular than in wildtypes and there were more often missing individual stereocilia in the shortest row in the mutants (Figure 10, A-H). In both inner and outer hair cells, stereocilia were arrayed in the cuticular plate in the normal V-shaped arrangement for OHCs or crescent for IHCs, and the angle of the V varied along the cochlear duct in a normal way in mutants, with a wider angle of the V in OHCs towards the base. Stereocilia were also arranged in rows of graded heights as usual, but with more irregularity in heights within each row in mutant bundles. All of these hair cell defects in IHCs and OHCs showed a gradient along the length of the cochlear duct with more severe defects towards the apex of the cochlea, but the abnormal IHC hair bundles extended further towards the base than abnormal OHC bundles. This tendency to more severe defects in the apex fits with the predominance of ABR threshold increases at low frequencies.

**Figure 9.**
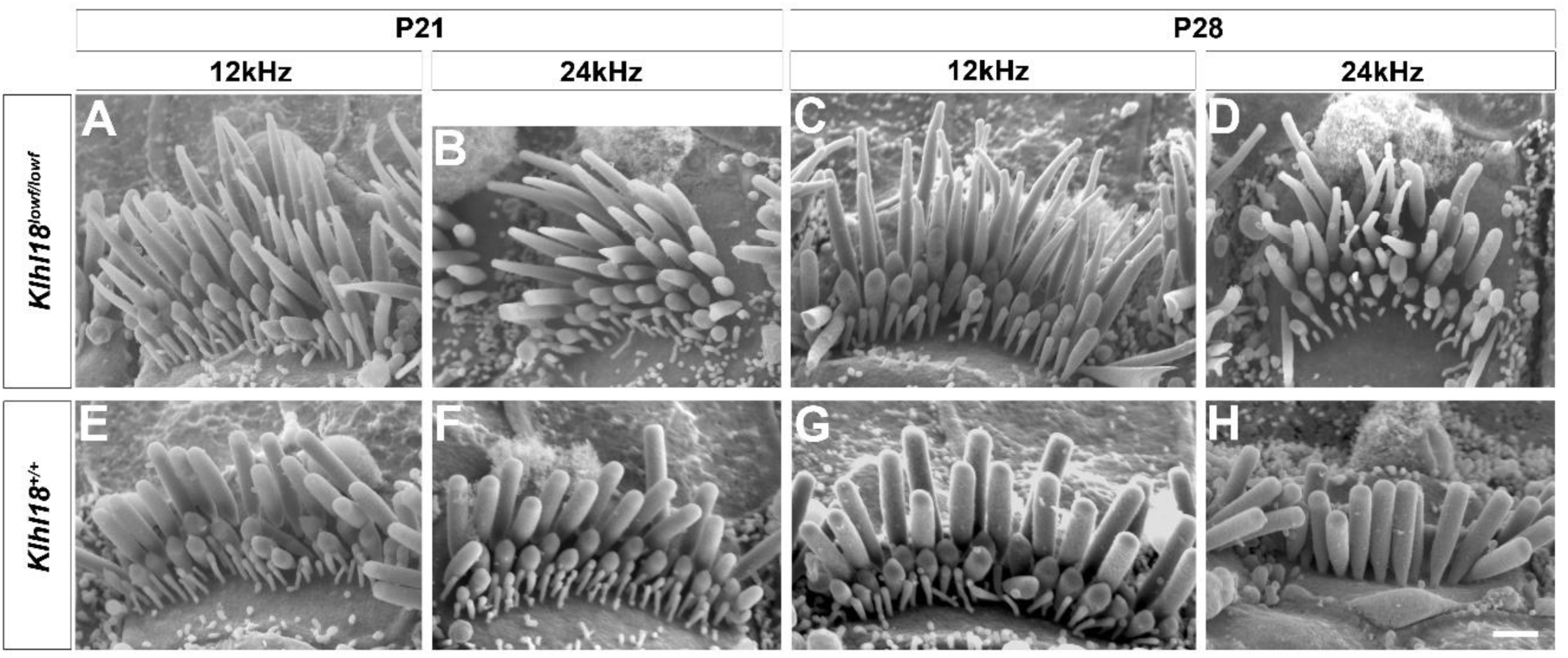
Scanning Electron Microscopy (SEM) of typical stereocilia bundles of Inner Hair Cells (IHC) in *Klhl18* mice. Representative images are shown from (**A-D**) *Klhl18^lowf/lowf^* homozygotes and (**E-H**) littermate wildtype mice. (**A, E**) P21 mice at the 12 kHz best-frequency region; (**B, F**) P21 mice at the 24 kHz best-frequency region; (**C, G**) P28 mice at the 12 kHz best-frequency region; (**D, H**) P28 mice at the 24 kHz best-frequency region. (**A-G**) show views from the modiolar side of the bundle, but (**H**) shows a view from the pillar side to illustrate the normal shape of the tallest row of stereocilia. Scale bar (on panel **H**) represents 1μm.

**Figure 10.**
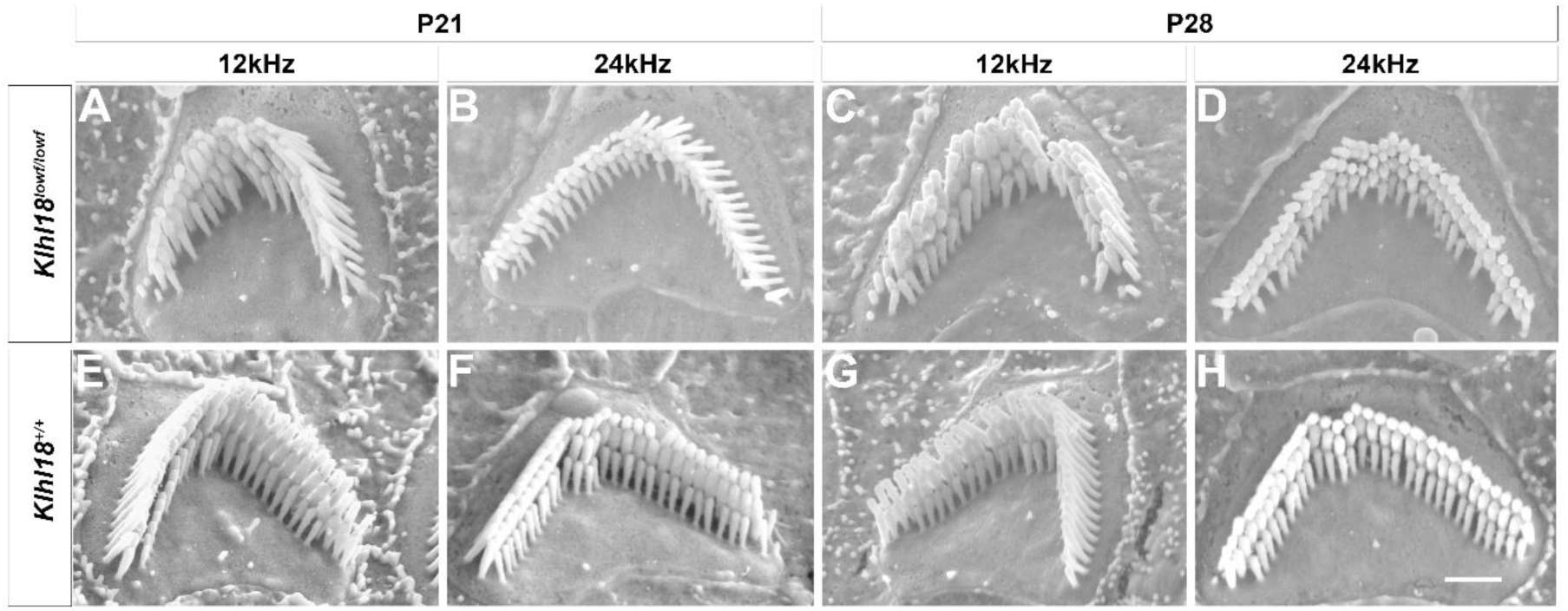
Scanning Electron Microscopy (SEM) of typical stereocilia bundles of Outer Hair Cells (OHC) in *Klhl18* mice. Representative images are shown from (**A-D**) *Klhl18^lowf/lowf^* homozygotes and (**E-H**) littermate wildtype mice. (**A, E**) P21 mice at the 12 kHz best-frequency region; (**B, F**) P21 mice at the 24 kHz best-frequency region; (**C, G**) P28 mice at the 12 kHz best-frequency region; (**D, H**) P28 mice at the 24 kHz best-frequency region. Scale bar (on panel **H**) represents 1μm.

Interestingly, we observed some minor tapering of IHC stereocilia in the apical 10-20% of the cochlear duct even in wildtypes. Heterozygotes showed an intermediate degree of IHC stereocilia tapering especially visible at P28 at the 3 and 6kHz best-frequency regions.

### Expression of *Klhl18* in organ of Corti supporting cells

We used the *Klhl18^tm1a^* allele to report the expected expression of *Klhl18* through the *LacZ* gene inserted into the allele (Figure 1B). Sections of cochleae from P13 heterozygotes revealed β-galactosidase activity, indicating *Klhl18* expression, in supporting cells of the organ of Corti (Hensen’s cells, Claudius’ cells, inner border cells) but not in the hair cells (Figure 11). The labelling was more pronounced in the apex compared with the basal turn and the staining distribution was consistently seen in all six heterozygotes examined. No expression of *Klhl18* was noted in hair cells, but it may be that expression levels at this age were below the levels detectable by the LacZ assay.

**Figure 11.**
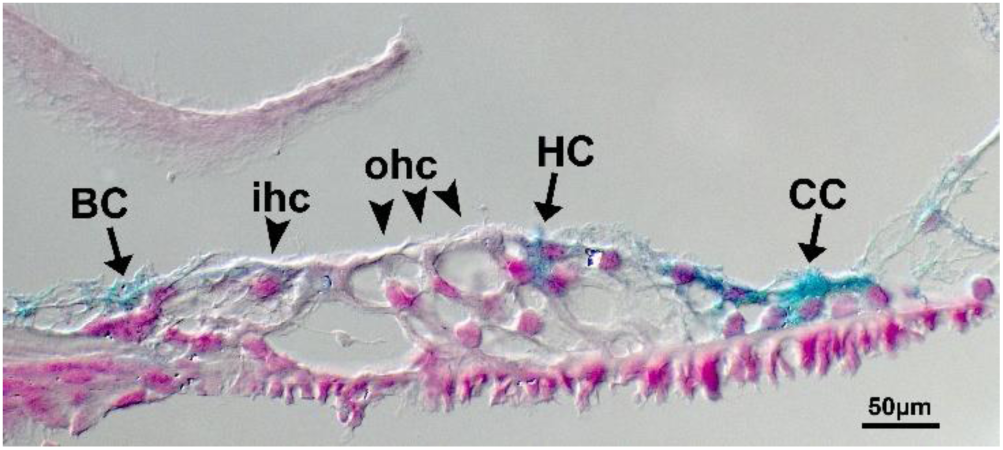
Expression of *Klhl18* in the organ of Corti. LacZ labelling of the organ of Corti from a P13 heterozygote for the *Klhl18^tm1a^* allele, showing expression (blue) in inner border cells (BS), Hensen’s cells (HC) and Claudius’ cells (CC), but not in either inner or outer hair cells (ihc, ohc). Nuclear Fast Red counterstaining is shown in pink.

## DISCUSSION

We have identified a key role for Klhl18 in the maintenance of cochlear function by tracking auditory responses from normal ABR thresholds at 3 weeks old to progressive deterioration of thresholds from 4 weeks onwards in mice with two different mutant alleles of the *Klhl18* gene. Low frequencies were predominantly affected, which is an unusual pattern. Outer hair cell function was normal judging from DPOAEs and CM responses, and the endocochlear potential was normal indicating normal stria vascularis function. Innervation of the organ of Corti appeared normal and IHCs had normal numbers of synapses with afferent neurons in mutants. However, scanning electron microscopy revealed abnormal tapering of IHC stereocilia in apical regions of the cochlea responsive to lower frequency tones, with a more normal appearance of stereocilia in the basal, high frequency regions.

There are relatively few cases where hearing impairment affects mainly the lower frequency ranges, and these have a variety of underlying pathologies. Mice lacking beta-tectorin have a disrupted tectorial membrane resulting in elevated CAP thresholds at low frequencies (Russell et al., 2007). The headbanger mutation of *Myo7a* results in defective stereocilia bundles, resulting in low frequency hearing impairment (Rhodes et al., 2004). Other examples include a low frequency sensorineural hearing impairment in a mouse model of Muenke Syndrome (Mansour et al 2009) and a low frequency hearing loss of unknown pathology in *Camsap3* mutant mice (Ingham et al 2019). In humans, low frequency hearing loss is rare and can be associated with non-syndromic deafness DFNA1 (Lynch et al., 1997), Wolfram Syndrome 1 (WFS1, Bespalova et al., 2001; Cryns et al., 2003; Sun et al., 2011) and with Meniere’s Disease (Lopez-Escamez et al., 2015). The inclusion of Klhl18 into this short list represents an important finding.

Stereocilia tapering and disorganised bundles were observed not only at 4 weeks old, where the apical preponderance of tapering correlated with reduced sensitivity to low frequency sounds, but also at 3 weeks old when ABR thresholds were normal in the *Klhl18^lowf^* mutants. Furthermore, less extensive tapering and hair bundle disorganisation was found in the 3 and 6 kHz best-frequency regions of heterozygotes and wildtypes. This was a surprising finding, reported by all three observers who were blinded to genotype. What might explain this? As the tapering phenotype is seen mainly in the longest row of stereocilia, it may be that sufficient tip-links survive to produce normal IHC signal transduction and hearing function. By 4 weeks of age, the bundle disruption and tapering effect is more extensive, possibly with the consequence that any remaining tip-links can no longer support normal transduction, causing hearing deterioration. Another potential explanation is that the frequency-place map of the cochlea may be shifted in mice from this colony (in both mutants and wildtypes) allowing hair cells from more basal regions to respond to lower frequencies than usually expected. Our observations suggest that an IHC stereocilia defect is an additional cochlear pathology that can be hidden behind normal ABR thresholds, a phenomenon previously linked to auditory nerve myelination defects (Wan and Corfas, 2017), to cochlear synaptopathy (Kujawa and Liberman 2009) and to auditory nerve dysfunction (Reijntjes et al 2019).

*Klhl18* is a member of the Kelch-like family of proteins, with domains that interact with many other proteins, including interaction of Kelch repeats with actin (Adams et al., 2000). Deficiency of Klhl18 may therefore result in a range of cellular problems in highly actin-rich cells such as IHCs, and in particular may affect the actin core of stereocilia leading to the tapering phenotype we observed. The tapering stereocilia of IHCs in *Klhl18* mutant mice is similar to that noted in *Espn^+/je^* mutant mice by Sekerkova et al. (2011). These heterozygote jerker mutant mice showed a transient tapering of apical turn (80% of cochlear length) IHC stereocilia which is apparent at P5 but not at P10. In more extreme apical regions (>95%), this tapering persists until at least 8 months of age. However, they did not observe tapering in their wildtype mice, but these were CBA/CaJ inbred strain control mice and not littermates of the *Espn^+/je^* mice (Sekerkova et al., 2011). Missing stereocilia from the shortest row in OHCs was also reported in the jerker heterozygotes, similar to our finding in *Klhl18* mutants. Espin is known to be involved in actin bundling. Loss of other actin-interacting proteins can also produce occasional narrowed IHC stereocilia, for example plastin1 (Taylor et al., 2015) and Eps8L2 (Furness et al 2013).

The related family member Klhl19 is known to co-localize with Myo7a in IHCs in the inner ear (Velichkova et al., 2002). Mutations in *Myo7a* are associated with stereocilia defects (Self et al. 1998) including tapering IHC stereocilia (Prosser et al. 2008) and low frequency hearing impairment (Rhodes et al. 2004). If, like Klhl19, Klhl18 is able to bind Myo7a, it may play a role in stereocilia stability and maintenance in a similar way.

The observation of tapering stereocilia even in wildtypes is unusual and may indicate that there is another genetic variant in this colony that can also influence stereocilia formation. The mice in this study were all homozygous for a targeted mutation of *Mab21l4* (also known as *2310007B03Rik*) but this mutation alone does not cause increased ABR thresholds (Ingham et al., 2019). Other mutations may have occurred spontaneously and become fixed within the current colony. Whatever any role of the *Mab21l4* mutation might be, the *Klhl18^lowf^* mutation must exacerbate the tapering phenotype as homozygotes are the most severely affected.

We found abnormalities in the SP measured from the round window in response to low frequency tones. The SP is a complex potential reflecting hair cell depolarisation, with contributions from both OHCs and IHCs (Harvey and Steel 1992; Pappa et al., 2019). However, as all other observations of OHCs in this study suggest they are functioning normally in the *Klhl18* mutant mice, the abnormalities noted in the SP are most likely due to IHC dysfunction. The hair bundle defect may impair the IHC receptor potential, disrupting normal voltage changes in the IHC that are required for normal activation of Ca^2+^ channels (Ohn et al., 2016) and therefore synaptic transmission (Brandt et al., 2003).

Our finding of *Klhl18* expression in organ of Corti supporting cells but not in hair cells was surprising, given that the main structural defect we saw was in the hair bundles of hair cells. Supporting cells play important roles in establishing the regular arrangement of hair cells during development (e.g. Wan et al., 2013), but the type of stereocilia defects we observed in *Klhl18* mutants has not been reported previously in cases where the gene is predominantly expressed in supporting cells rather than in hair cells. Single cell RNA-Seq data (Kolla et al. 2020) suggests that *Klhl18* is indeed expressed in both IHC and OHCs at P0, but we examined expression at P13. This suggests that Klhl18 may be required in early hair cell development only.

A recent report from Chaya et al. (2019) demonstrated that *Klhl18* is expressed in mouse photoreceptors and that *Klhl18^tm1b^* homozygous mutants had impaired electroretinographic responses associated with mislocalization of rod transducin α subunit. The Cul3-Klhl18 ubiquitin ligase was found to target Unc119, promoting its degradation and thus modulating transducin α translocation between rod inner and outer segments during light adaptation. Klhl18 is also involved in ubiquitylation of Aurora-A (Moghe et al., 2012). However, there is no clear link between the role of Klhl18 as a ubiquitin ligase and the stereocilia defects that we observed.

Auditory Neuropathy Spectrum Disorder (ANSD) is a complex and heterogeneous condition in humans. ANSD can be relatively easy to diagnose, characterised by normal otoacoustic emissions along with abnormal ABRs, but there are a wide range of underlying causes, genetic or otherwise (eg Moser and Starr 2016). Our findings suggest that the *KLHL18* gene will be a good candidate for involvement in human ANSD. Further studies of the hearing impairment associated with mutation of *Klhl18* will contribute to a wider understanding of pathophysiological mechanisms contributing to low frequency hearing impairment and to ANSD.

## Acknowledgements

This work was supported by Wellcome (098051, 100699, and WT089622MA), the Medical Research Council (G0300212 to KPS), Royal National Institute for Deaf People (to KPS and NJI) and the EC (MSCA-ITN-2016-LISTEN-722098 to KPS). We thank the Wellcome Sanger Institute Mouse Genetics Project for access to the *Klhl18^tm1a(KOMP)Wtsi^* and *Mab21l4^tm1a(KOMP)Wtsi^* mutant mice; Dr Morag A. Lewis for genotyping advice and colony management; Dr Elisa Martelletti for advice on confocal microscopy; Dr Selina A. Pearson for ABR recordings to help to establish the *Klhl18^lowf^* colony; and Dr Douglas C. Fitzpatrick for helpful suggestions on round window recordings. The funders had no role in study design, data collection and analysis, decision to publish, or preparation of the manuscript.

## EXTENDED DATA FIGURES

**EXTENDED DATA Figure 2-1.**
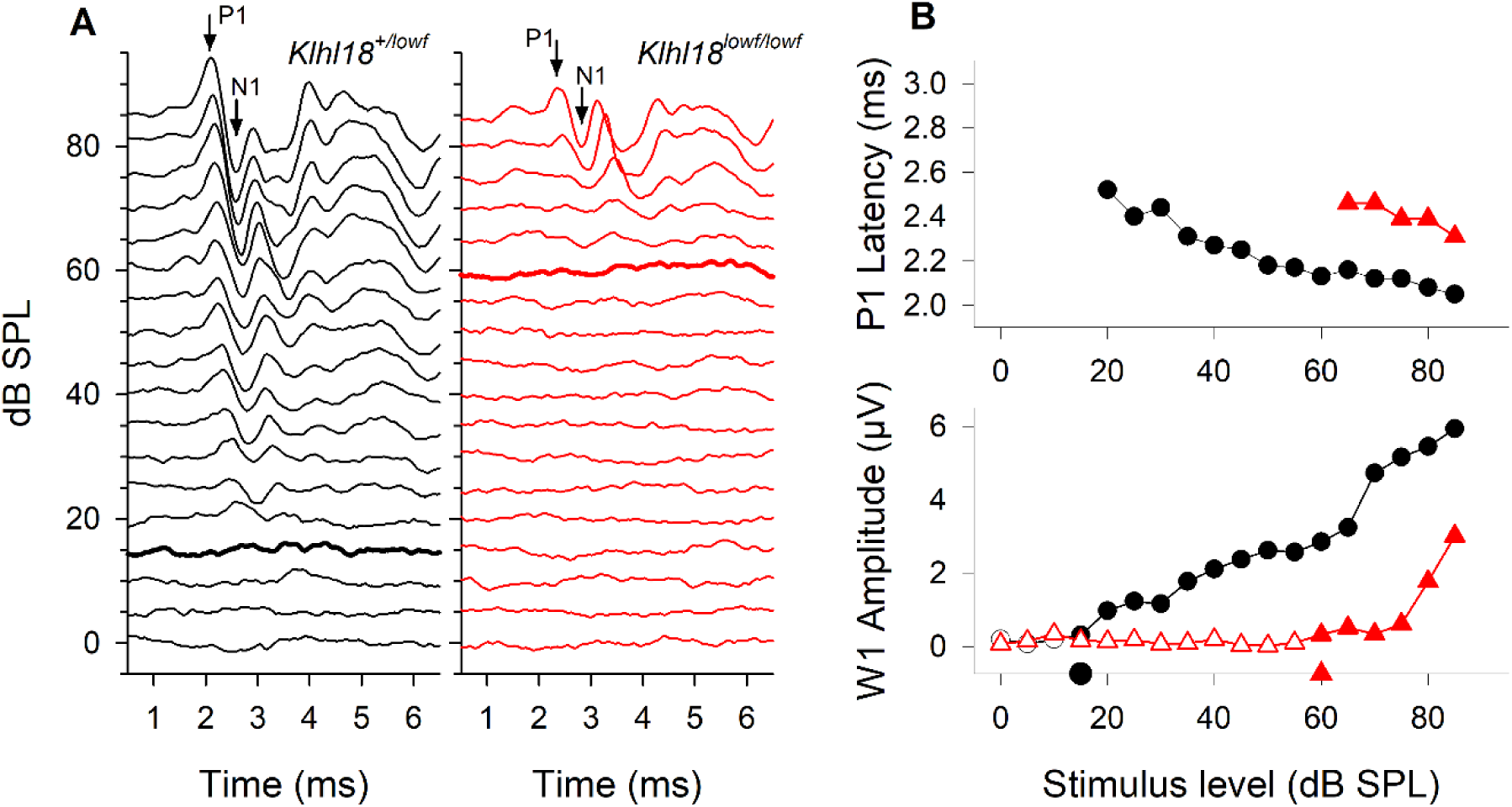
**Recordings of Auditory Brainstem Responses (ABRs) in *Klhl18^lowf^* mice. A**, Representative ABRs recorded in response to 12 kHz tone pips presented from 0-85 dB SPL are plotted for a *Klhl18^+/lowf^* control mouse (black) and a *Klhl18^lowf/lowf^* mutant mouse (red). P1 and N1 indicate the positions of the first positive and negative peaks of the waveform, from which amplitude and latency measurements are make. The thickened line on each stack of ABR traces indicates the visually-determined threshold level for these responses (15 dB SPL for the control mouse; 60 dB SPL for the mutant mouse). **B**, Input-Output functions for ABR wave 1 amplitude and P1 latency measurements (from the traces shown in A) are plotted as a function of stimulus sound pressure level (dB SPL), for the control mouse (black circles) and mutant mouse (red up-triangles). Sub-threshold points are indicated by open symbols. Supra-threshold points are indicated by filled symbols. The estimated threshold for each mouse is indicated by the appropriate single symbol on the abscissa of the wave 1 amplitude function.

**EXTENDED DATA Figure 2-2.**
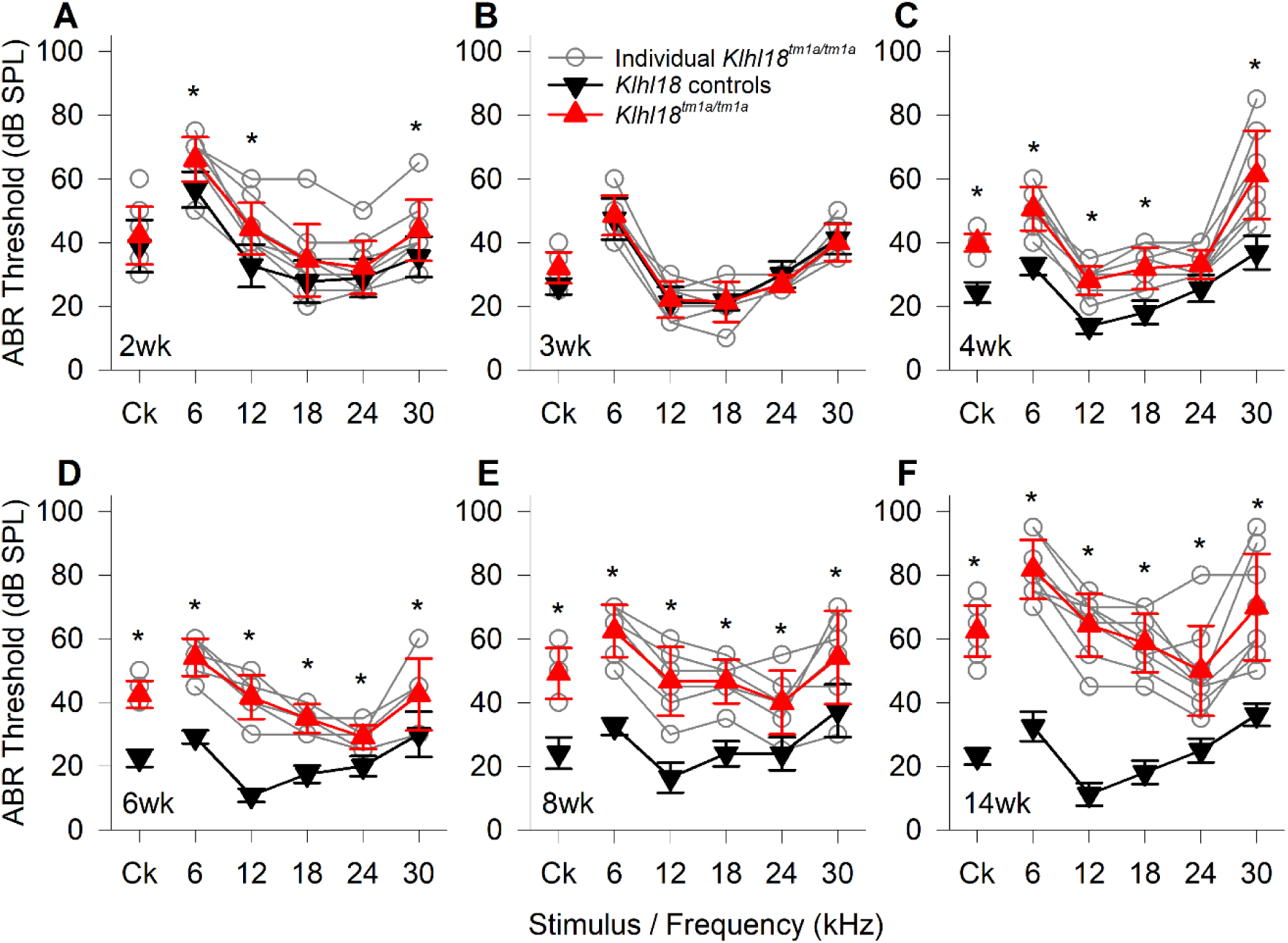
Age-Related changes in ABR thresholds of *Klhl18^tm1a^* mice. A summary of ABR thresholds estimated from control (wildtype or heterozygous) and mutant (homozygous) are shown for *Klhl18^tm1a^* mice aged 2 weeks (**A**, n=9 control, including 3 wildtype, n=9 mutant), 3 weeks (**B**, n=4 control, all wildtype, n=7 mutant), 4 weeks (**C**, n=8 control, including 3 wildtype, n=8 mutant), 6 weeks (**D**, n=6 control, including 1 wildtype, n=6 mutant), 8 weeks (**E**, n=8 control, including 1 wildtype, n=8 mutant) and 14 weeks (**F**, n=8 control, including 3 wildtype, n=8 mutant). Thresholds from individual mutant mice are indicated by grey circles. Mean threshold (±SD) are plotted for control mice as black down-triangles and for mutant mice as red up-triangles. Ck : Click stimulus. A mixed-effects model statistical analysis between control and mutant thresholds gave the following results; 2 weeks, F (1, 48) = 39.659, p=8.790x10^-8^; 3 weeks, F (1, 54) = 0.253, p=0.617; 4 weeks, F (1, 84) = 171.196, p=1.000 x10^-15^; 6 weeks, F (1, 30) = 245.412, p=1.000 x10^-15^; 8 weeks, F (1, 30) = 265.386, p=1.000 x10^- 15^; 14 weeks, F (1, 84) = 523.580, p=1.000 x10^-15^. Sidak’s multiple comparisons test was used to examine the difference between control and mutant thresholds for each stimulus. Significant differences are given in Table 1-1 and indicated here by *.

**EXTENDED DATA Figure 2-3.**
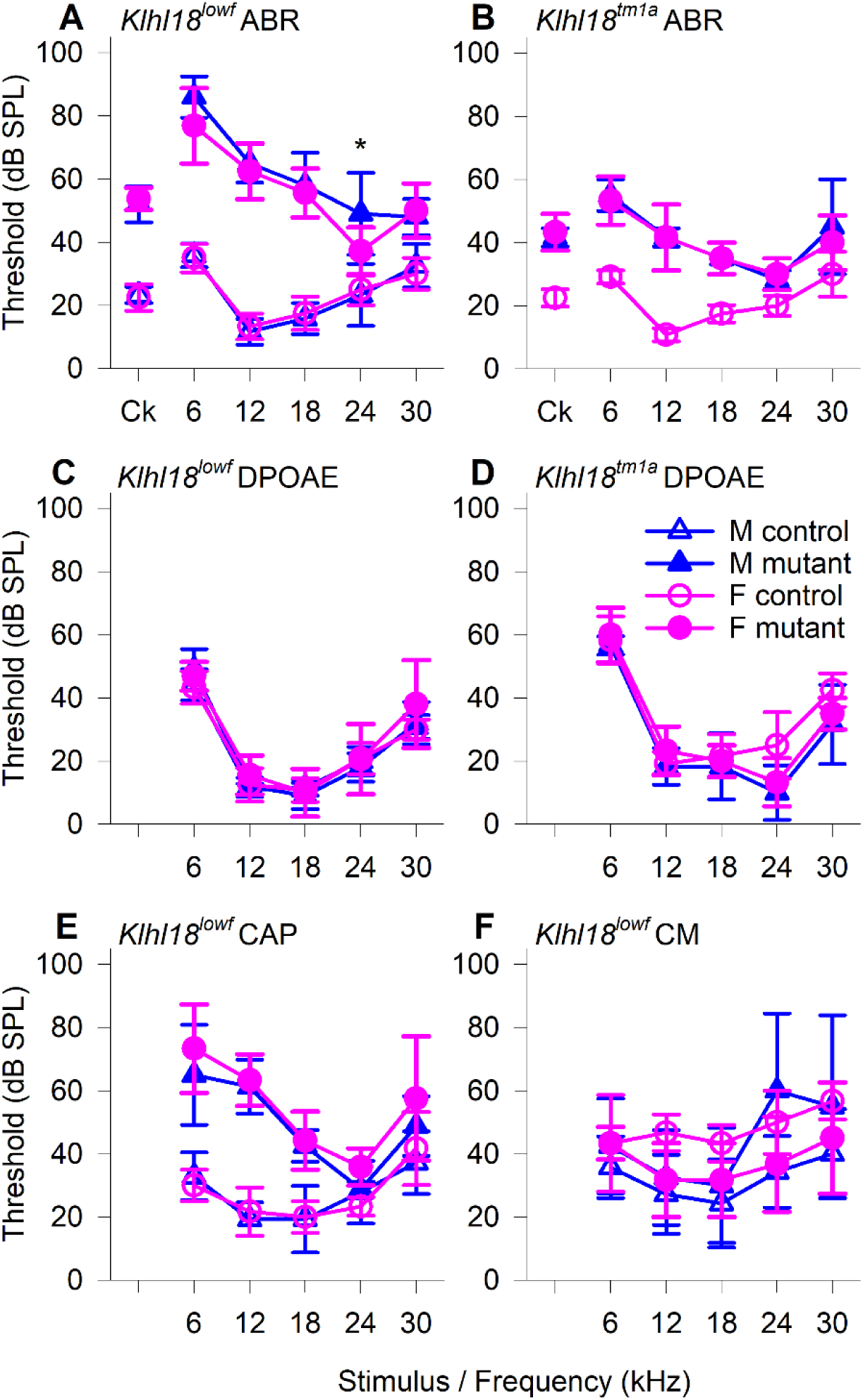
Male vs Female responses of 6 weeks old *Klhl18* mice. Mean (± SD) thresholds for male (blue triangles) and female (pink circles) Klhl18 control mice (open symbols) and Klhl18 mutant mice (filled symbols) are plotted in **A** ABR thresholds of *Klhl18^lowf^* mice (n=6 control male, n=6 control female, n=5 mutant male and n=8 mutant female), **B** ABR thresholds of *Klhl18^tm1a^* mice(n=6 control female, n=3 mutant male and n=3 mutant female), **C** DPOAE thresholds of *Klhl18^lowf^* mice(n=6 control male, n=6 control female, n=5 mutant male and n=8 mutant female), **D** DPOAE thresholds of *Klhl18^tm1a^* mice(n=6 control female, n=3 mutant male and n=3 mutant female), **E** CAP thresholds of *Klhl18^lowf^* mice (n=7 wildtype male, n=3 wildtype female, n=4 mutant male and n=6 mutant female) and **F** CM thresholds of *Klhl18^lowf^* mice (n=7 wildtype male, n=3 wildtype female, n=4 mutant male and n=6 mutant female). A mixed-effects model statistical analysis between male and female control and mutant thresholds gave the following results; ABR thresholds for *Klhl18^lowf^* control mice, F (1, 58) = 0.012, p=0.914; ABR thresholds for *Klhl18^lowf^* mutant mice, F (1, 7) = 1.700, p=0.234; ABR thresholds for *Klhl18^tm1a^* control mice (n/a, no male mice tested); ABR thresholds for *Klhl18^tm1a^* mutant mice, F (1, 2) = 0.0816, p=0.802; DPOAE thresholds for *Klhl18^lowf^* control mice, F (1, 5) = 0.000, p=1.000; DPOAE thresholds for *Klhl18^lowf^* mutant mice, F (1, 7) = 0.470, p=0.515; DPOAE thresholds for *Klhl18^tm1a^* control mice (n/a, no male mice tested); DPOAE thresholds for *Klhl18^tm1a^* mutant mice, F (1, 2) = 100.000, p=9.852 x10^-3^; CAP thresholds for *Klhl18^lowf^* control mice, F (1, 6) = 0.013, p=0.911; CAP thresholds for *Klhl18^lowf^* mutant mice, F (1, 5) = 2.737, p=0.159; CM thresholds for *Klhl18^lowf^* control mice, F (1, 6) = 4.513. p=0.078; CM thresholds for *Klhl18^lowf^* mutant mice, F (1, 5) = 0.411, p=0.550. Sidak’s multiple comparisons test was used to examine the difference between control and mutant thresholds for each stimulus. Significant differences are given in extended data Table 1-1 and indicated here by *.

**EXTENDED DATA Figure 3-1.**
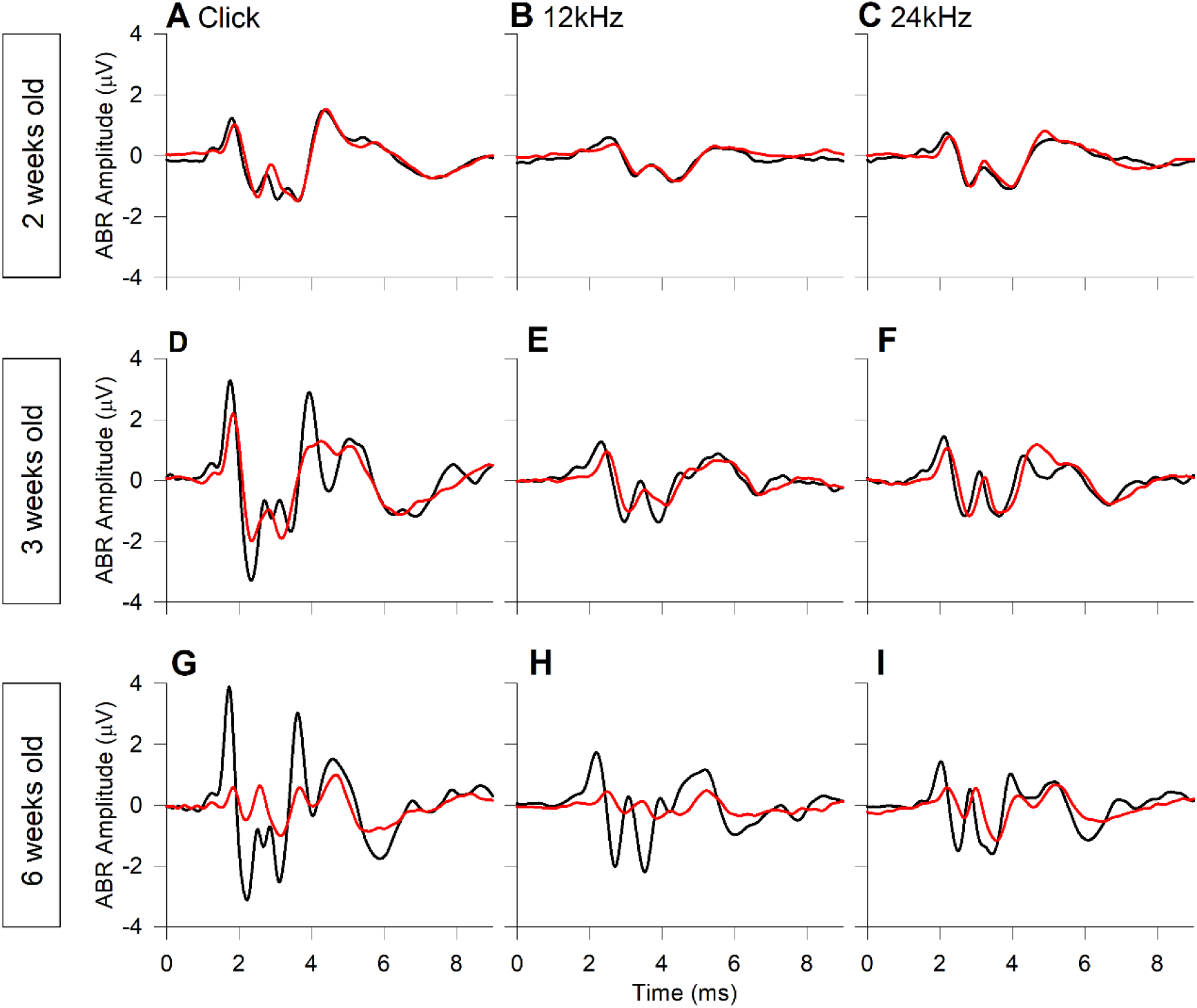
Mean ABR waveforms of *Klhl18^tm1a^* mice at 65 dB SPL. Group averaged ABR waveforms, evoked by 65 dB SPL stimuli, are plotted for mice aged 2 weeks (**A-C**, n=9 control, including 3 wildtype, n=9 mutant), 3 weeks (**D-F**, n=4 control, all wildtype, n=7 mutant), 6 weeks (**G-I**, n=6 control, including 1 wildtype, n=6 mutant), in response to click stimuli (A,D,G), 12 kHz stimuli (**B,E,H**) and 24 kHz stimuli (**C,F,I**). Mean amplitude waveforms are plotted for the same wildtype mice (black lines) and mutant mice (red lines) described in figure 2-2.

**EXTENDED DATA Figure 3-2.**
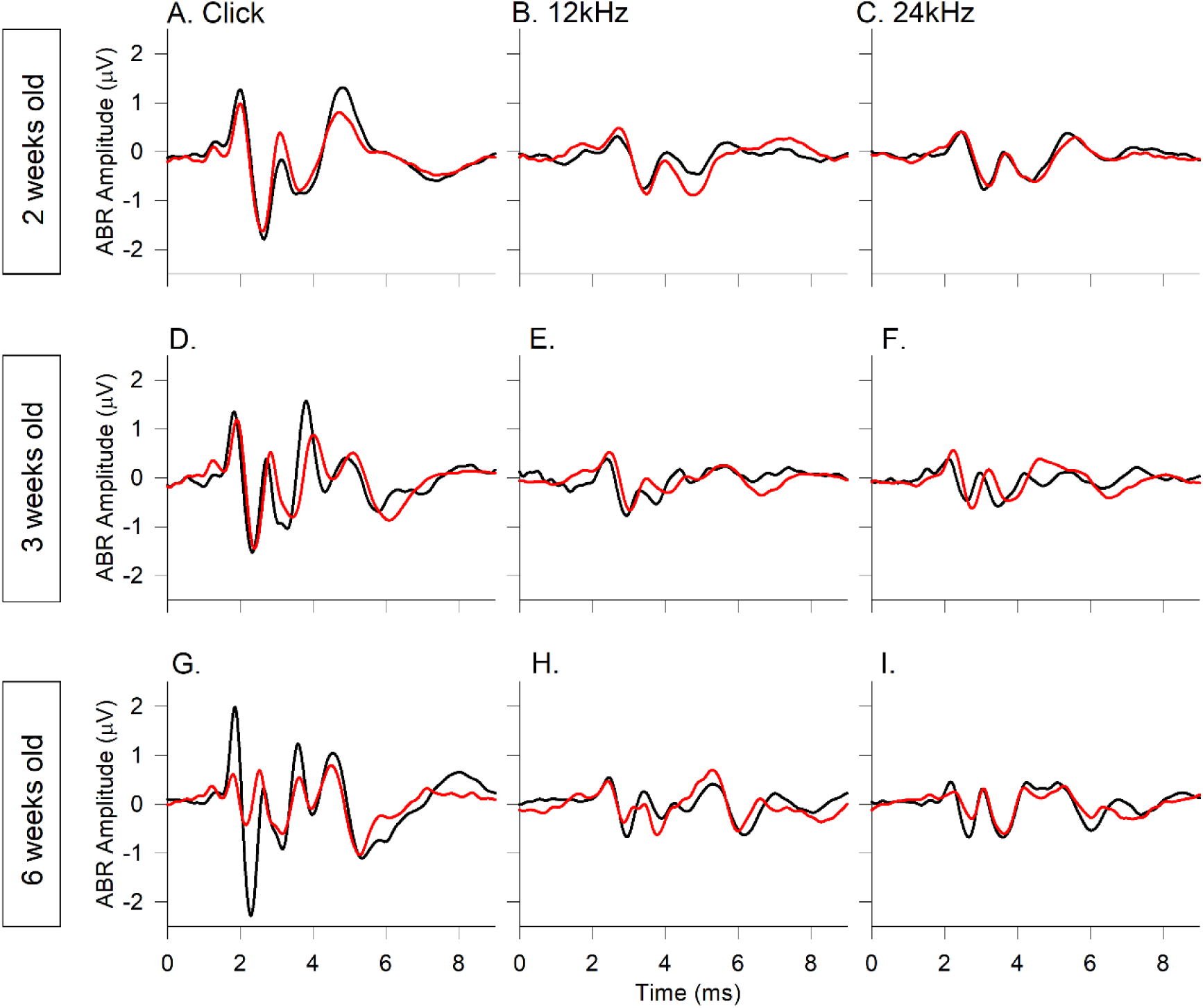
Mean ABR waveforms of *Klhl18^lowf^* mice at 20 dB SL. Group averaged ABR waveforms, evoked by stimuli presented at 20 dB above threshold (20 dB sensation level, dB SL), are plotted for mice aged 2 weeks (**A-C**, n=13 control, n=12 mutant), 3 weeks (**D-F**, n=8 control, n=19 mutant), 6 weeks (**G-I**, n=17 control, n=10 mutant), in response to click stimuli (**A,D,G**), 12 kHz stimuli (**B,E,H**) and 24 kHz stimuli (**C,F,I**). Mean amplitude waveforms are plotted for the same wildtype mice (black lines) and mutant mice (red lines) described in figure 2.

**EXTENDED DATA Figure 3-3.**
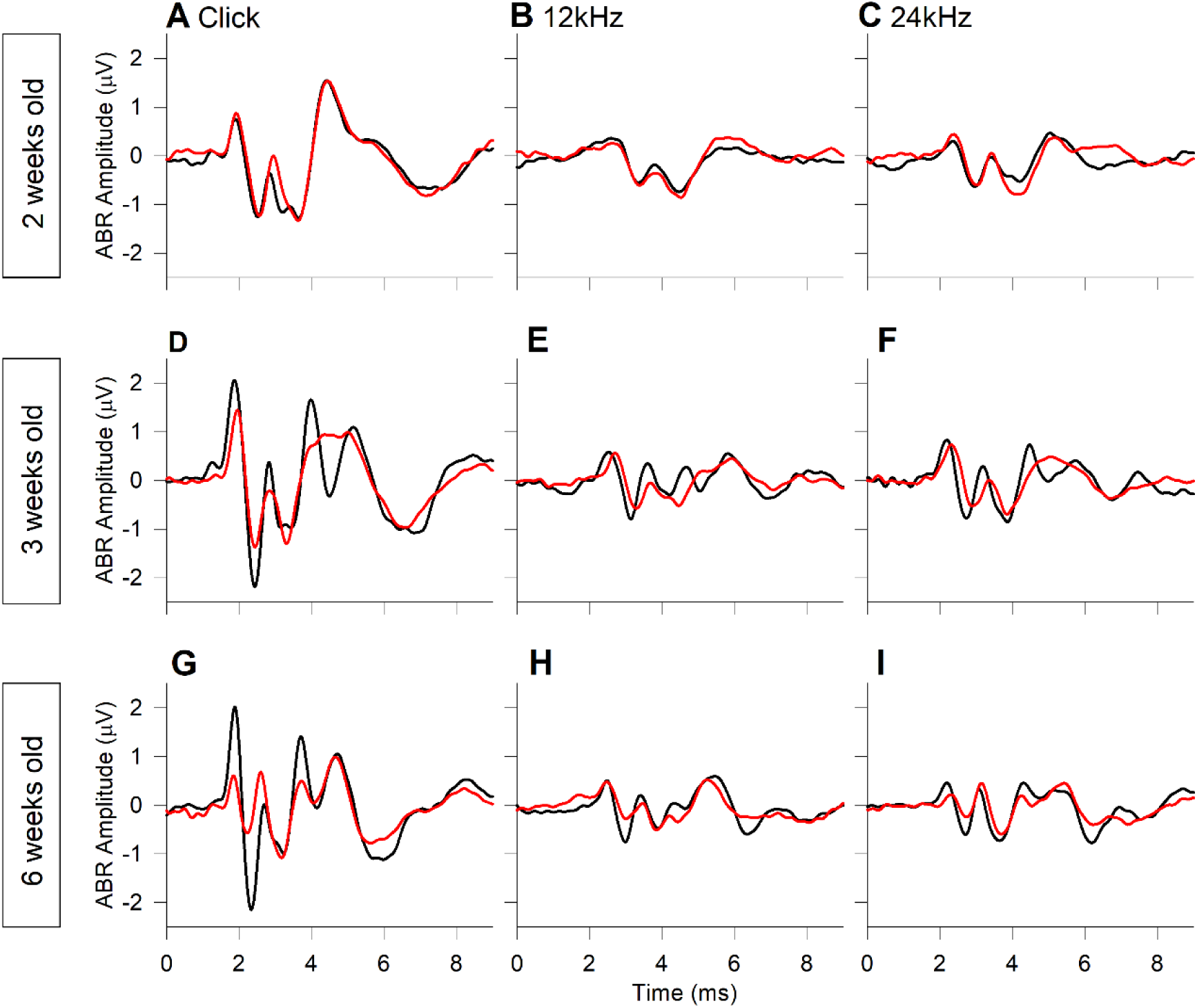
Mean ABR waveforms of *Klhl18^tm1a^* mice at 20 dB SL. Group averaged ABR waveforms, evoked by stimuli presented at 20 dB above threshold (20 dB sensation level, dB SL), are plotted for mice aged 2 weeks (**A-C**, n=9 control, including 3 wildtype, n=9 mutant), 3 weeks (**D-F**, n=4 control, all wildtype, n=7 mutant), 6 weeks (**G-I**, n=6 control, including 1 wildtype, n=6 mutant), in response to click stimuli (**A,D,G**), 12 kHz stimuli (**B,E,H**) and 24 kHz stimuli (**C,F,I**). Mean amplitude waveforms are plotted for the same wildtype mice (black lines) and mutant mice (red lines) described in figure 2-2.

**EXTENDED DATA Figure 4-1.**
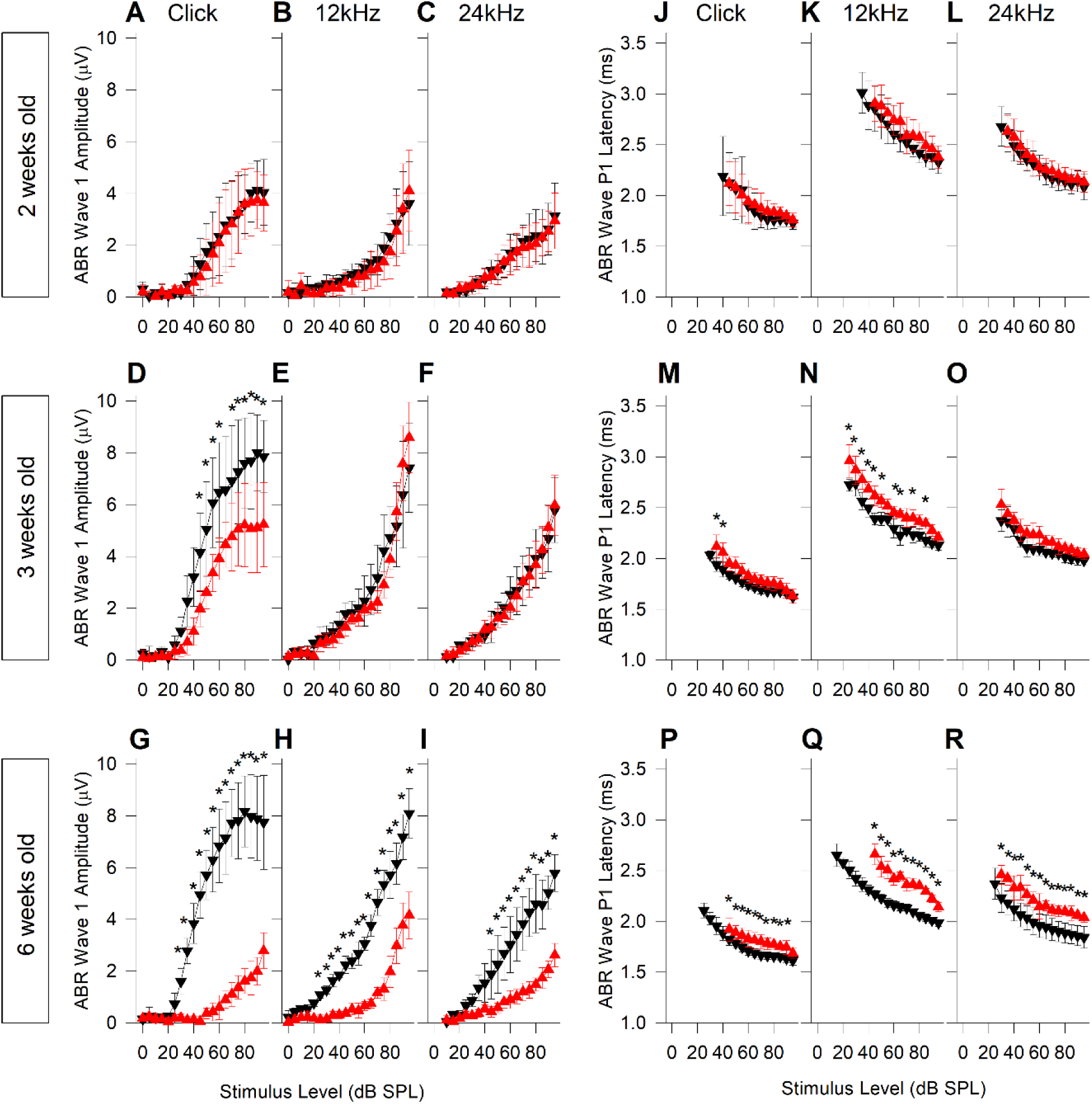
ABR wave 1 amplitude and latency input-output functions in *Klhl18^tm1a^* mice. Mean ABR wave 1 P1-N1 peak-to-peak amplitude (±SD) as a function of stimulus level is plotted for mice aged 2, 3 and 6 weeks old are plotted in **A-C** (n=9 controls, including 3 wildtypes; n=9 mutants), **D-F** (n=4 wildtype controls, n=7 mutants) and **G-I** (n=6 controls, including 1 wildtype; n=6 mutants), respectively. Results are plotted for click stimuli (**A,D,G**), 12 kHz tones (**B,E,H**) and 24 kHz tones (**C,F,I**). Data from control mice are plotted as black down-triangles. Data from mutant mice are plotted as red up-triangles. Mean ABR wave P1 latency (±SD) as a function of stimulus level is plotted for the same mice aged 2, 3 and 6 weeks old are plotted in **J-L**, **M-O** and **P-R**, respectively. Results are plotted for click stimuli (**J,M,P**), 12kHz tones (**K,N,Q**) and 24kHz tones (**L,O,R**). A mixed-effects model statistical analysis between control and mutant thresholds gave the following results; Wave 1 amplitude evoked by clicks at 2 weeks, F (1, 278) = 2.139, p=0.145; 3 weeks, F (1, 180) = 103.491, p=1.000 x10^-15^; 6 weeks, F (1, 200) = 1195.900, p=1.000 x10^-15^; Wave 1 amplitude evoked by 12 kHz at 2 weeks, F (1, 268) = 3.499, p=0.062; 3 weeks, F (1, 174) = 3.402, p=0.067; 6 weeks, F (1, 196) = 1210.838, p=1.000 x10^-15^; Wave 1 amplitude evoked by 24 kHz at 2 weeks, F (1, 260) = 0.931, p=0.336; 3 weeks, F (1, 160) = 0.340, p=0.561; 6 weeks, F (1, 160) = 370.953, p=1.000 x10^-15^; P1 latency evoked by clicks at 2 weeks, F (1, 176) = 2.398, p=0.123; 3 weeks, F (1, 117) = 57.494, p=8.822 x10^-12^; 6 weeks, F (1, 55) = 149.479, p=1.000 x10^-15^; P1 latency evoked by 12 kHz at 2 weeks, F (1, 176) = 23.864, p=2.312 x10^-6^; 3 weeks, F (1, 45) = 234.216, p=1.000 x10^-15^; 6 weeks, F (1, 55) = 1698.126, p=1.000 x10^-15^; P1 latency evoked by 24 kHz at 2 weeks, F (1, 208) = 8.333, p=4.304 x10^-3^; 3 weeks, F (1, 126) = 54.587, p=1.000 x10^-15^; 6 weeks, F (1, 70) = 206.472, p=1.000 x10^-15^. Sidak’s multiple comparisons test was used to examine the difference between control and mutant data for each stimulus. Significant differences are given in Table 1-1 and indicated here by *.

**EXTENDED DATA Figure 4-2.**
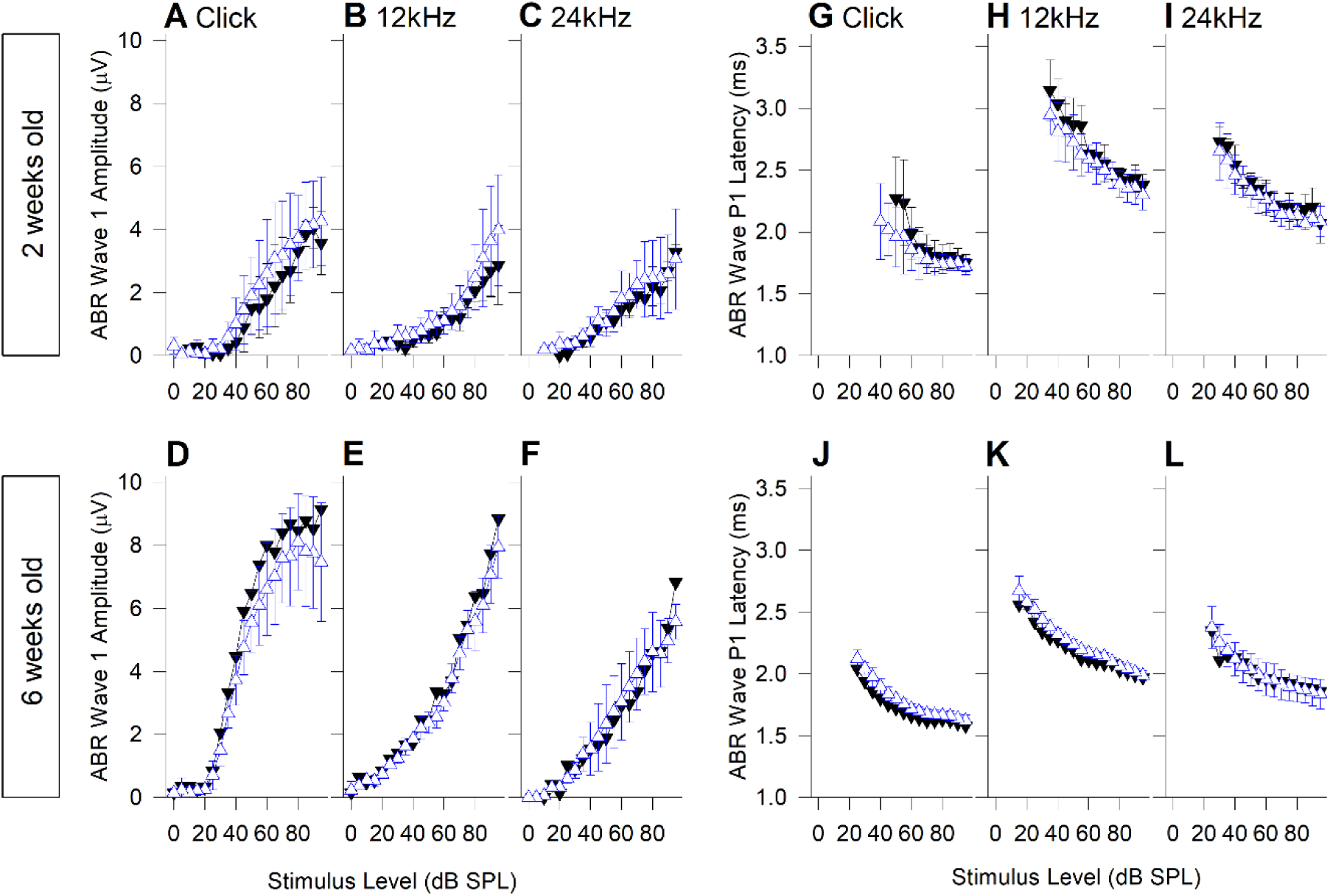
ABR wave 1 amplitude and latency input-output functions in *Klhl18^tm1a^* heterozygote mice. Mean ABR wave 1 P1-N1 peak-to-peak amplitude (±SD) as a function of stimulus level is plotted for mice aged 2 and 6 weeks old in **A-C** (n=3 wildtype; n=6 heterozygotes), and **D-F** (n=1 wildtype; n=5 heterozygotes), respectively. Results are plotted for click stimuli (**A,D**), 12 kHz tones (**B,E**) and 24 kHz tones (**C,F**). Data from wildtype mice are plotted as black down-triangles. Data from heterozygote mice are plotted as open blue up-triangles. Mean ABR wave P1 latency (±SD) as a function of stimulus level is plotted for the same mice aged 2 and 6 weeks old are plotted in **G-I** and **J-L**, respectively. Results are plotted for click stimuli (**G,J**), 12 kHz tones (**H,K**) and 24 kHz tones (**I,L**). A mixed-effects model statistical analysis between control and mutant thresholds gave the following results; Wave 1 amplitude evoked by clicks at 2 weeks, F (1, 32) = 16.188, p=3.279 x10^-4^; Wave 1 amplitude evoked by 12 kHz at 2 weeks, F (1, 30) = 7.244, p=1.152 x10^-2^; Wave 1 amplitude evoked by 24 kHz at 2 weeks, F (1, 30) = 3.616, p=0.067; P1 latency evoked by clicks at 2 weeks, F (1, 70) = 7.899, p=6.408 x10^-3^; P1 latency evoked by 12 kHz at 2 weeks, F (1, 26) = 23.682, p=4.781 x10^-5^; P1 latency evoked by 24 kHz at 2 weeks, F (1, 28) = 11.159, p=2.380 x10^-3^. Sidak’s multiple comparisons test was used to examine the difference between control and mutant data for each stimulus. Significant differences are given in Table 1-1 and indicated here by *.

**EXTENDED DATA Figure 5-1.**
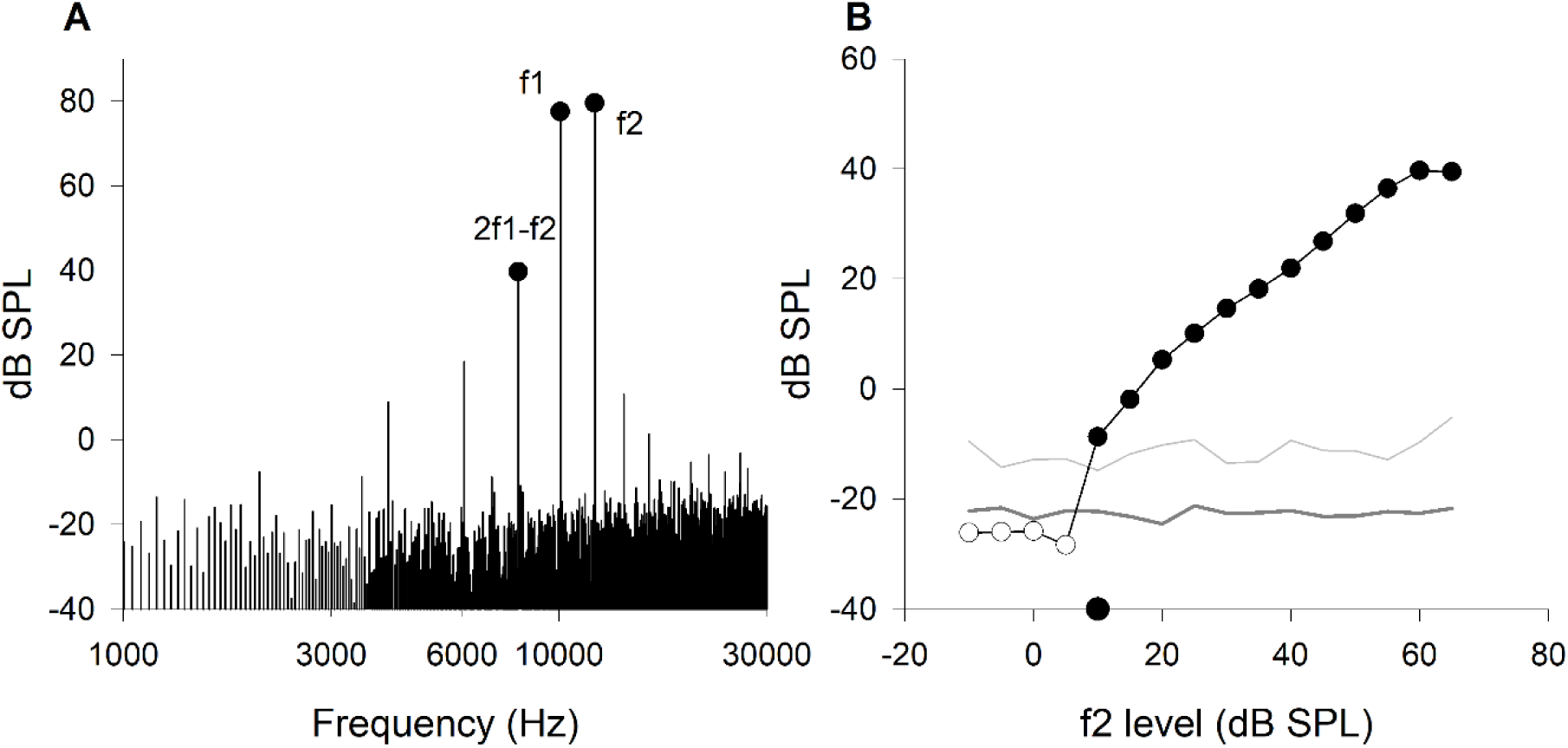
**Recordings of Distortion Product Otoacoustic Emissions (DPOAEs) in *Klhl18^lowf^* mice**. **A**. A representative DPOAE frequency spectrum is plotted for a response recorded with a stimulus of 2 frequency components, f1 (10000 Hz, 70 dB SPL) and f2 (12000 Hz, 60 dB SPL), for a *Klhl18^+/+^* mouse. The spectral peaks of the two stimulus tones are labelled f1 and f2. The measured DPOAE is labelled 2f1-f2. These peaks are further highlighted by filled circles. Similar spectra were recorded using f2 levels from -10 dB to 65 dB and used to create a growth function for the 2f1-f2 DPOAE. **B**. For the same *Klhl18^+/+^* mouse, DPOAE amplitude is plotted as a function of f2 level (circles). The thick grey line indicates the mean noise-floor amplitude for each response. The thin grey line indicates 2 standard deviations (SD) above the mean for the noise-floor. This line is used to estimate a threshold for the DPOAE component; the lowest f2 stimulus level where the DPOAE is greater than 2 SDs above the noise-floor. Sub-threshold DPOAEs are indicated by open circles. Supra-threshold DPOAEs are indicated by black filled circles. The estimated threshold is indicated by the single symbol on the abscissa.

**EXTENDED DATA Figure 5-2.**
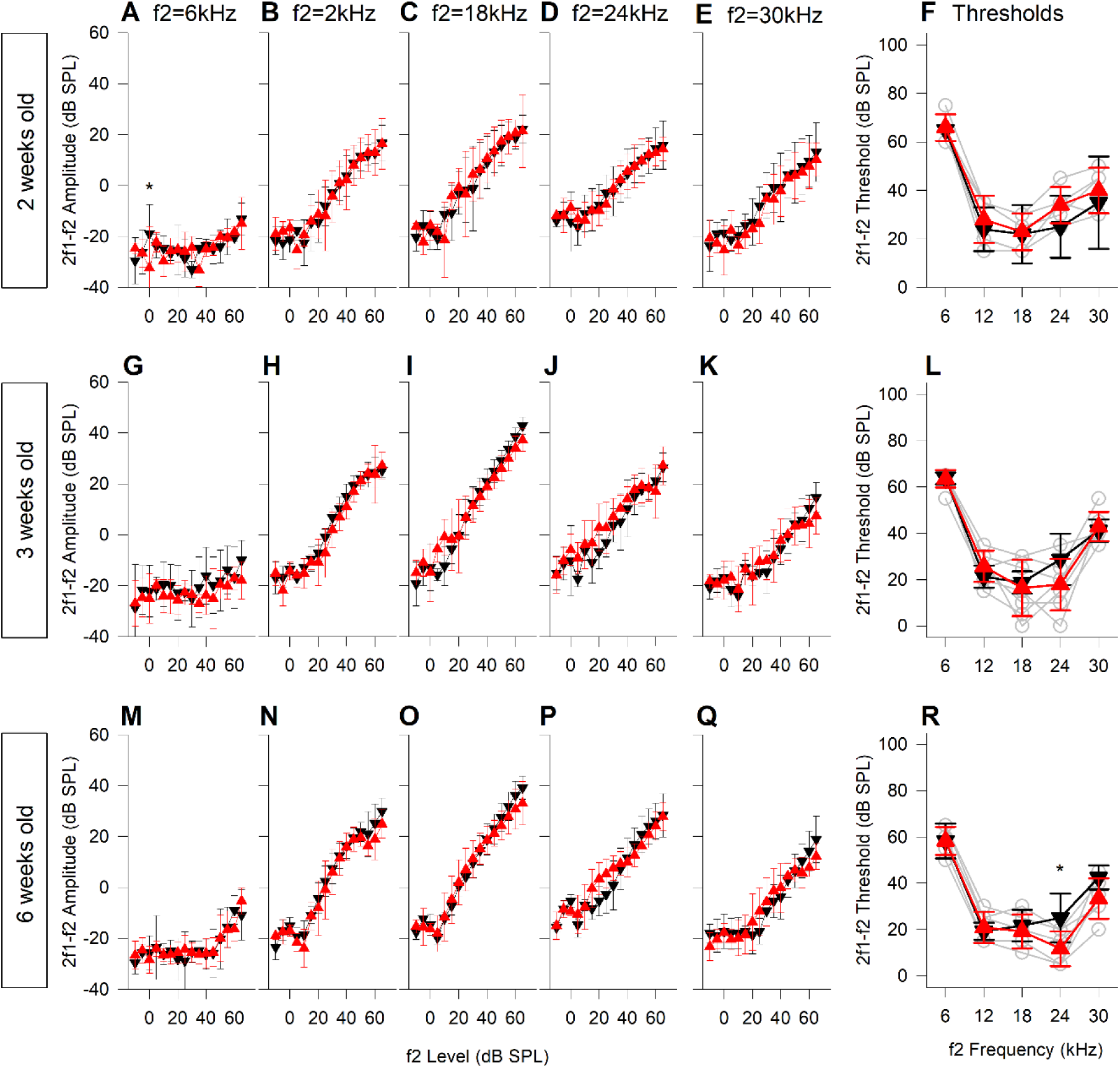
DPOAE growth and thresholds in *Klhl18^tm1a^* mice. DPOAE results from mice aged 2, 3 and 6 weeks old are plotted in **A-F** (n=5 controls, n=5 mutants), **G-L** (n=4 controls, n=7 mutants) and **M-N** (n=6 controls, n=6 mutants), respectively. Data from control mice are plotted as black down-triangles. Data from mutant mice are plotted as red up-triangles. The mean (±SD) amplitude of the 2f1-f2 DPOAE (dB SPL) is plotted as a function of f2 stimulus level (dB SPL) for f2 frequencies of 6 kHz (**A, G, M**), 12 kHz (**B, H, N**), 18 kHz (**C, I, O**), 24 kHz (**D, J, P**) and 30 kHz (**E, K, Q**). Mean threshold (±SD) of the 2f1-f2 DPOAE (derived from individual growth functions, eg shown in Figure 5-1) and plotted in **F**, **L** and **R** for mice aged 2 weeks, 3 weeks and 6 weeks respectively. In addition to the mean data, thresholds from individual mutant mice are plotted as grey open circles. A mixed-effects model statistical analysis between control and mutant data gave the following results; DPOAE amplitude evoked by an f2 of 6 kHz at 2 weeks, F (1, 112) = 0.121, p=0.729; 3 weeks, F (1, 144) = 5.802, p=1.727 x10^-2^; 6 weeks, F (1, 160) = 0.025, p=0.875; DPOAE amplitude evoked by an f2 of 12 kHz at 2 weeks, F (1, 48) = 0.052, p=0.821; 3 weeks, F (1, 144) = 4.751, p=3.090 x10^-2^; 6 weeks, F (1, 80) = 6.850, p=1.060 x10^-2^; DPOAE amplitude evoked by an f2 of 18 kHz at 2 weeks, F (1, 48) = 0.016, p=0.901; 3 weeks, F (1, 144) = 0.078, p=0.781; 6 weeks, F (1, 80) = 1.038, p=0.311; DPOAE amplitude evoked by an f2 of 24 kHz at 2 weeks, F (1, 48) = 0.020, p=0.888; 3 weeks, F (1, 48) = 7.079, p=1.507 x10^-2^; 6 weeks, F (1, 80) = 1.770, p=0.187; DPOAE amplitude evoked by an f2 of 30 kHz at 2 weeks, F (1, 48) = 5.264, p=2.619 x10^-2^; 3 weeks, F (1, 144) = 0.044, p=0.834; 6 weeks, F (1, 80) = 0.116, p=0.0.735; DPOAE thresholds evoked across all stimulus frequencies, at weeks, F (1, 20) = 1.891, p=0.184; 3 weeks, F (1, 15) = 0.163, p=0.692; 6 weeks, F (1, 25) = 6.735, p=1.559 x10^-2^. Sidak’s multiple comparisons test was used to examine the difference between control and mutant data for each stimulus. Significant differences are given in Table 1-1 and indicated here by *.

**EXTENDED DATA Figure 6-1.**
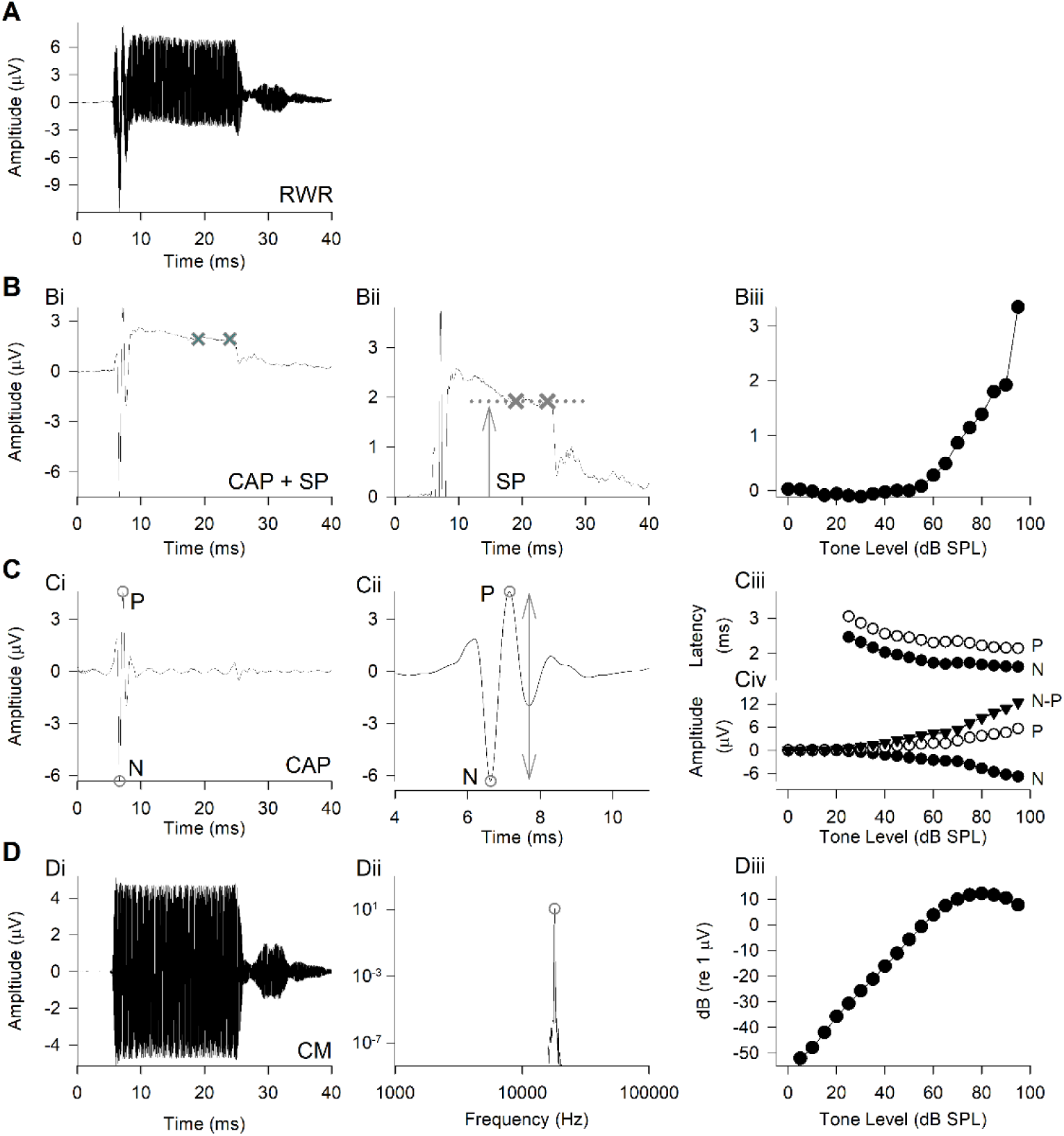
Round Window Response (RWR), analyses and parameters measured. **A.** A typical electrophysiological response recorded from the mouse round window with a 1 Hz-50 kHz bandwidth bioamplifier; in this case, in response to an 18 kHz tone pip at 90 dB SPL. **Bi.** The response in (A), low pass filtered at 3 kHz to remove the cochlear microphonic (CM) component and expose the summating potential (SP) and cochlear compound action potential (CAP). The “X” symbols indicate points on the response at 19 ms and 24 ms. Data points between these time values were averaged to generate the magnitude of the SP. **Bii** illustrates that the SP is measured as a voltage magnitude displaced from the zero baseline of the lowpass filtered response. The SP can be positive or negative relative to the baseline. **Biii.** SP is plotted against dB SPL to produce a SP input-out function (IOF). **Ci.** The RWR from (A), bandpass filtered (300 Hz – 3 kHz) to remove both the CM and SP components and isolate the CAP response, composed of a large negative peak (N) followed by a large positive peak (P), indicated by open circle symbols. **Cii** illustrates the same filtered response from Ci but with the abscissa expanded to clearly show the structure of the CAP waveform. Peaks N and P are again illustrated by open circles. The arrow indicates the peak-to-peak (N to P) magnitude of the CAP. The latency of the N and P peaks was also measured (corrected to account for the 5ms onset delay of the stimulus tone pip). **Ciii.** The latency of peaks N and P (filled and open circles respectively) are plotted against dB SPL to produce a latency IOF. **Civ.** Similarly, the amplitude of peaks N and P and the N-P amplitude are plotted against dB SPL to produce amplitude IOFs. **Di.** The RWR from (A) bandpass filtered, centred on the stimulus frequency with high- and low- pass corner frequencies at ± 100 Hz, to remove all components within the response other than the CM. **Dii.** The filtered CM response is limited to a 7-23 ms time window before applying a Fast Fourier Transformation; the resulting power spectrum is shown with the magnitude of the CM component indicated by an open circle. **Diii**. CM amplitudes measured across all stimulus level presented are plotted against dB SPL to produce a CM IOF.

**EXTENDED DATA Figure 6-2.**
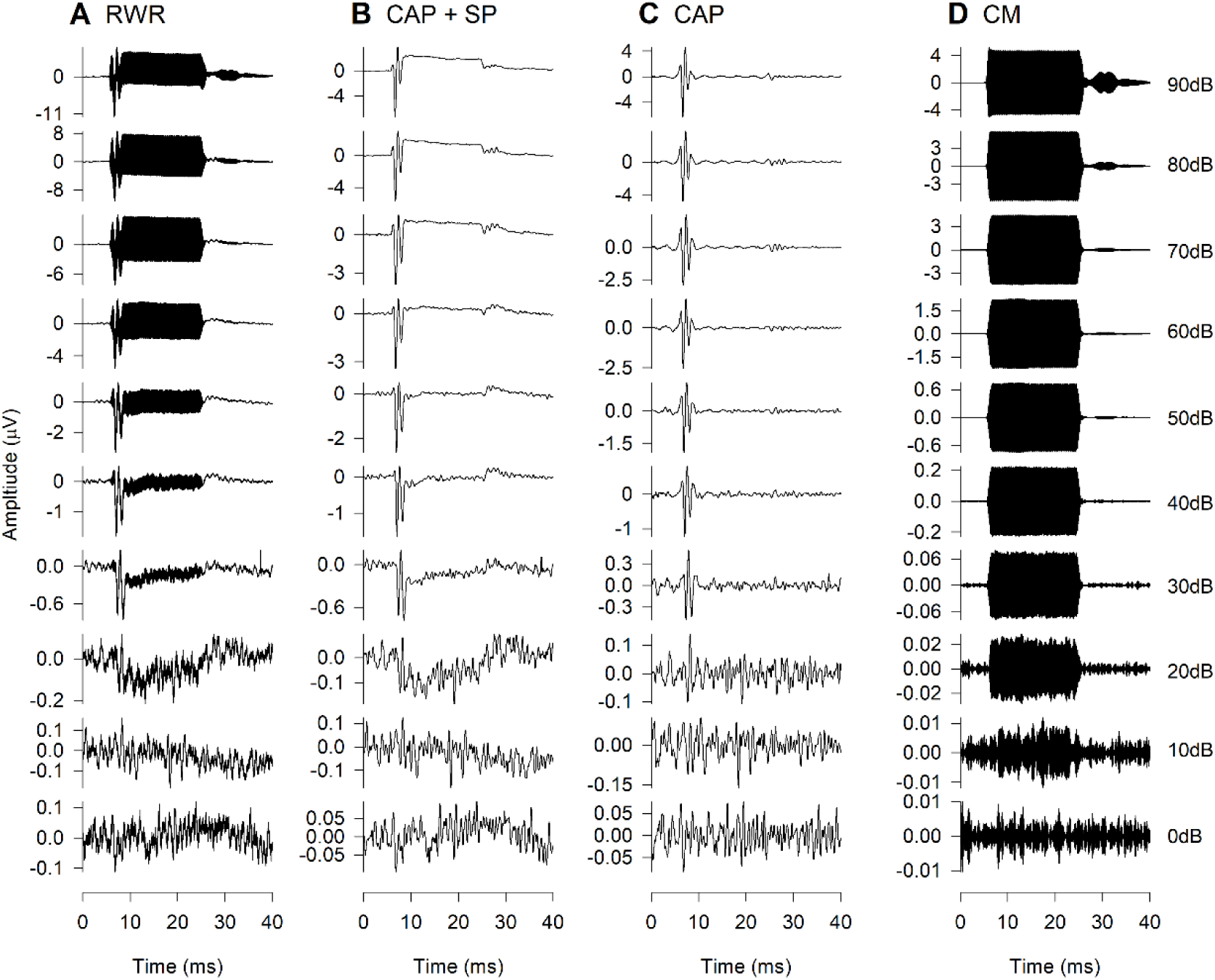
RWRs and components in a wildtype control mouse. Representative responses recorded for 18 kHz tones in a control mouse, over stimulus levels from 0 – 90 dB SPL are illustrated. **A.** Broadband RWRs (1 Hz – 50 kHz). **B.** Lowpass filtered (3 kHz) RWRs show responses containing the CAP and SP. **C.** Bandpass filtered (300 Hz - 3 kHz) RWRs show responses containing the CAPs. **D.** Bandpass filtered (18 kHz ± 100 Hz) RWRs show responses containing the CM.

**EXTENDED DATA Figure 6-3.**
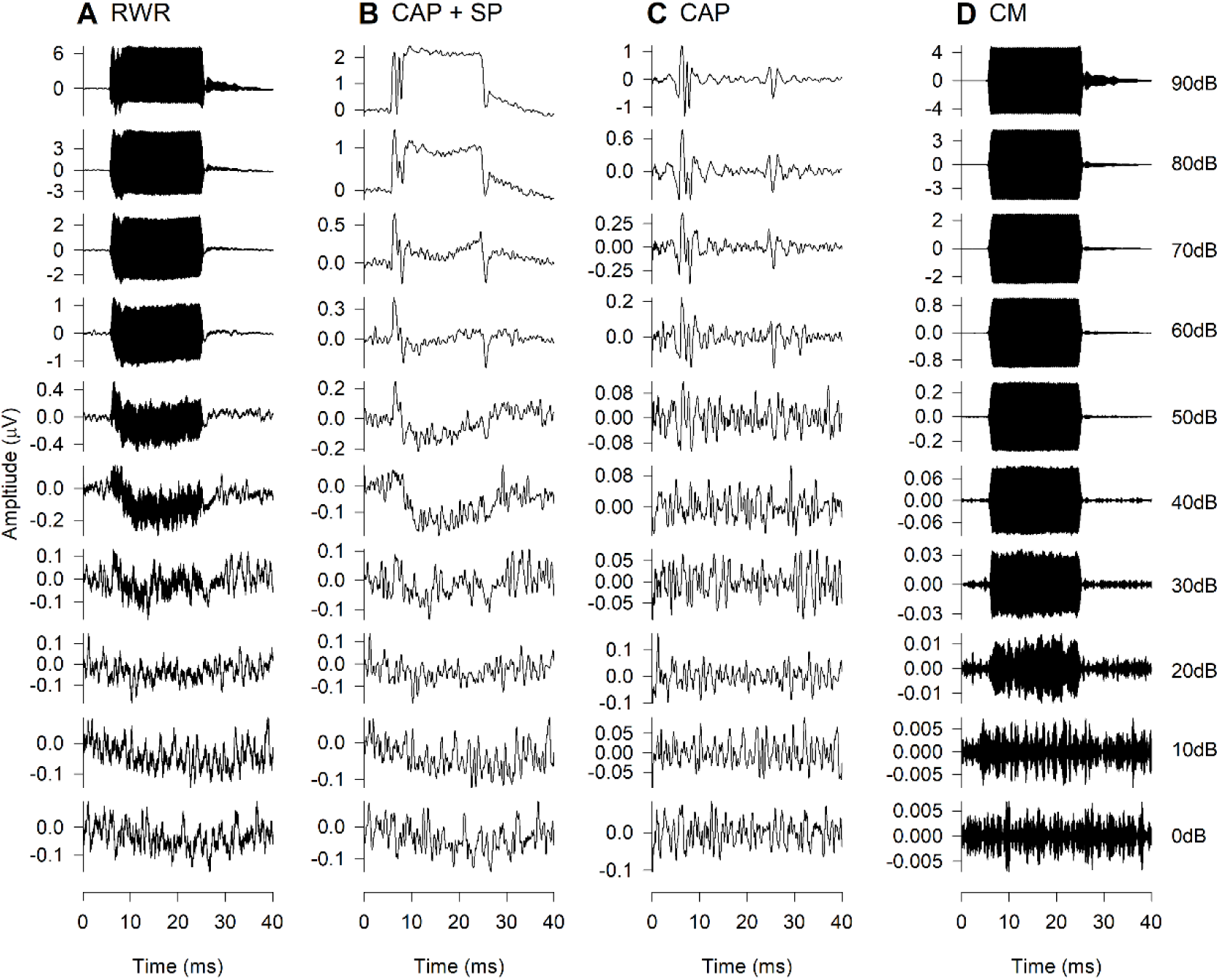
RWRs and components in a *Klhl18^lowf^* mutant mouse. Representative responses recorded for 18kHz tones in a control mouse, over stimulus levels from 0 – 90 dB SPL are illustrated. **A.** Broadband RWRs (1 Hz – 50 kHz). **B.** Lowpass filtered (3 kHz) RWRs show responses containing the CAP and SP. **C.** Bandpass filtered (300 Hz – 3 kHz) RWRs show responses containing the CAPs. **D.** Bandpass filtered (18 kHz ± 100 Hz) RWRs show responses containing the CM.

**EXTENDED DATA Figure 6-4.**
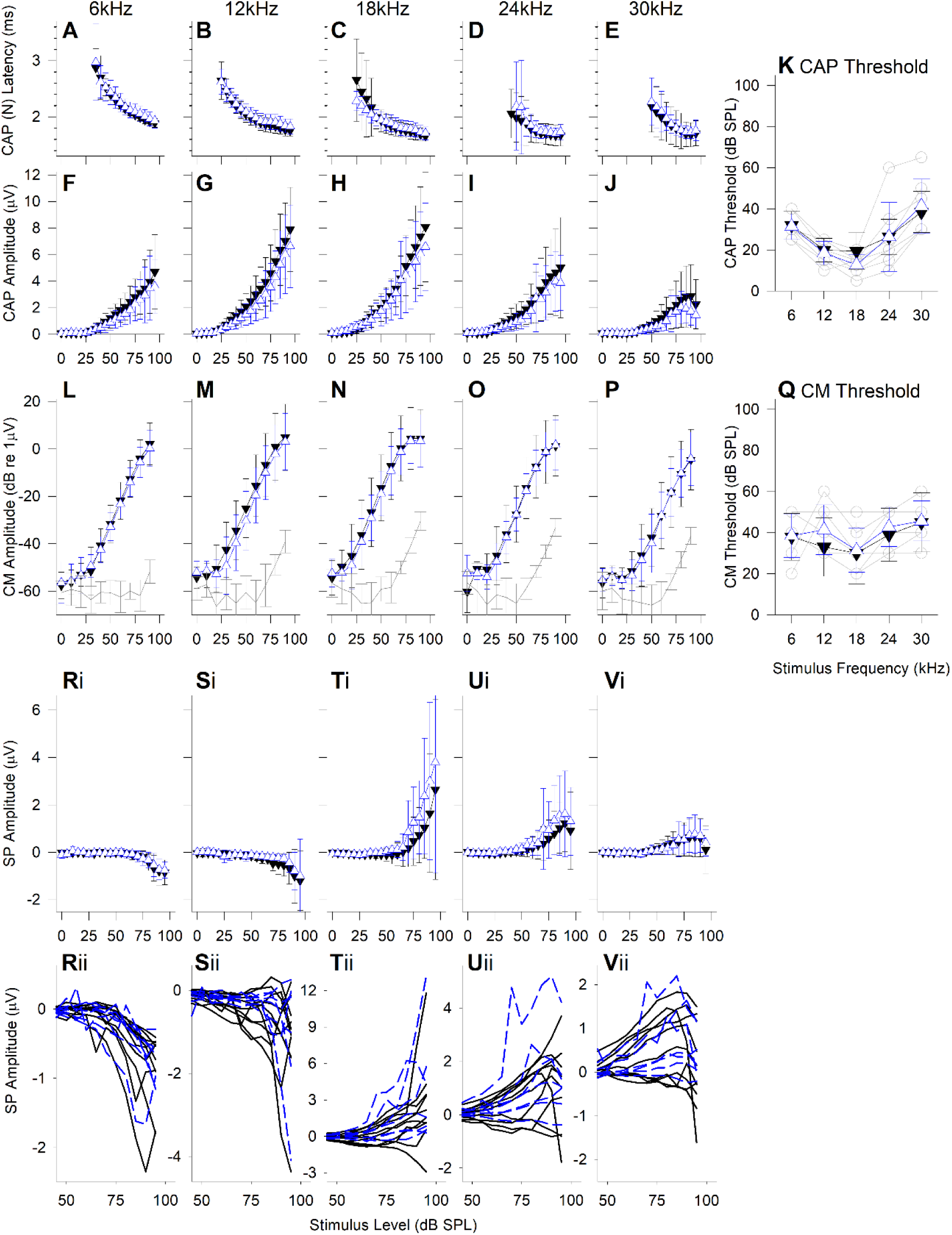
Round Window Response measurements from *Klhl18^+/lowf^* (heterozygote) mice. Compound Action Potential latency (of wave N) and amplitude (N-P amplitude) are plotted in **A**-**E** and **F**-**J**, respectively, for potentials measured in response to tones of 6, 12, 18, 24 and 30 kHz. Data are plotted as mean ± SD for wildtype control mice (n=10; black down-triangles) and heterozygote mice (n=7; red up-triangles). **K.** Mean ± SD of the CAP threshold is plotted against stimulus frequency for wildtype control mice (black down-triangles) and heterozygote mice (red up- triangles). Open circles and grey lines indicate CAP thresholds for individual heterozygote mice. **L**-**P** plot mean (± SD) Cochlear Microphonic amplitude (dB re 1 µV) against stimulus level (dB SPL) for wildtype mice (black down-triangles) and heterozygote mice (red up-triangles). The line plotted in grey represents the mean ± SD amplitude of stimulus artefact. **Q.** Mean ± SD of the estimated CM threshold is plotted against stimulus frequency for wildtype control mice (black down-triangles) and heterozygote mice (red up-triangles). Open circles and grey lines indicate CM thresholds for individual heterozygote mice. The grey line without symbols indicates the estimated magnitude of the stimulus artefact, as described in Figure 6. **Ri**-**Vi** plot mean (± SD) Summating Potential amplitude (µV) against stimulus level (dB SPL) for wildtype mice (black down-triangles) and heterozygote mice (red up-triangles). **Rii**-**Vii** plot SP IOFs for individual control (black lines) and mutant (red lines) mice.

**EXTENDED DATA TABLE 1-1.**
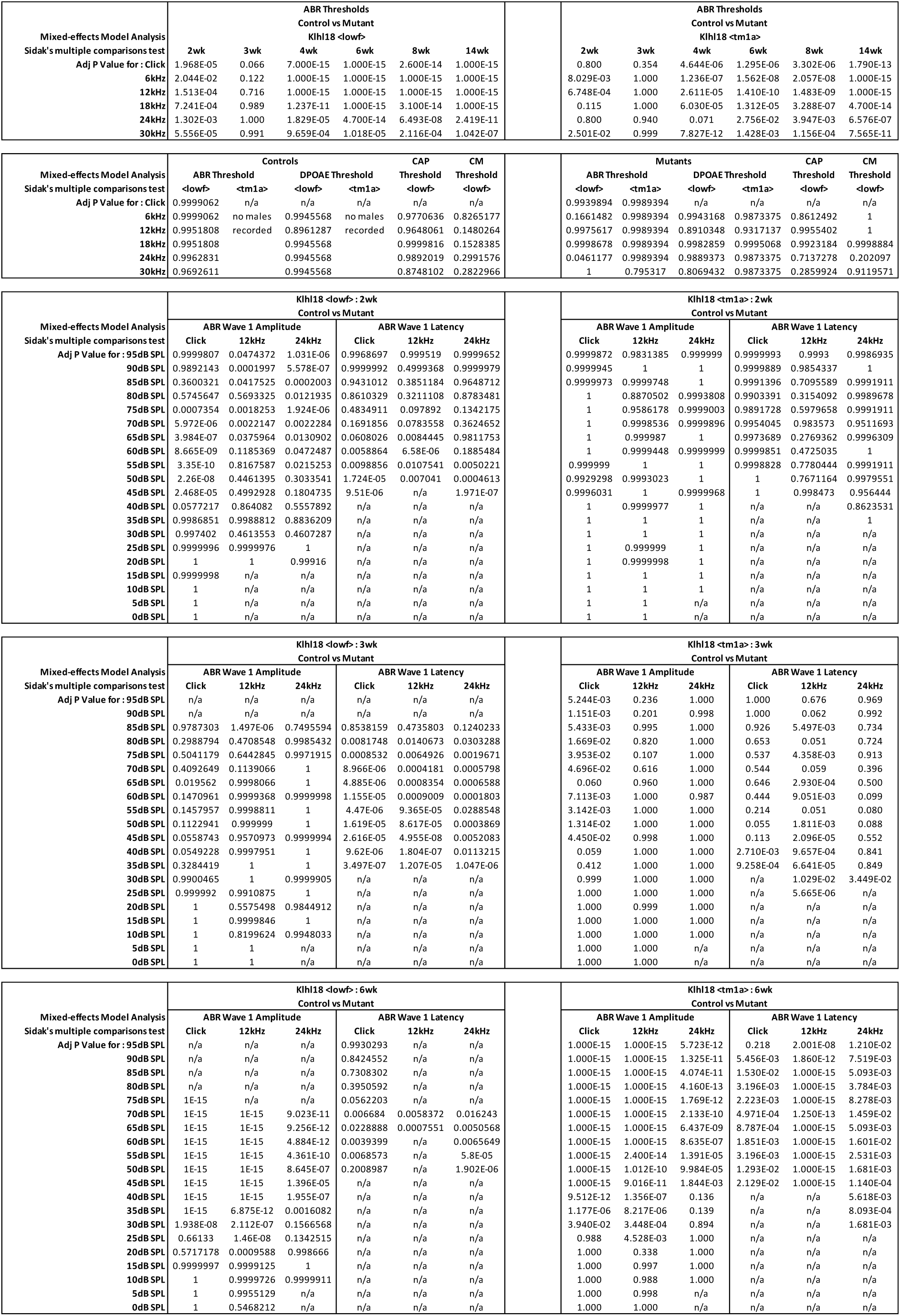
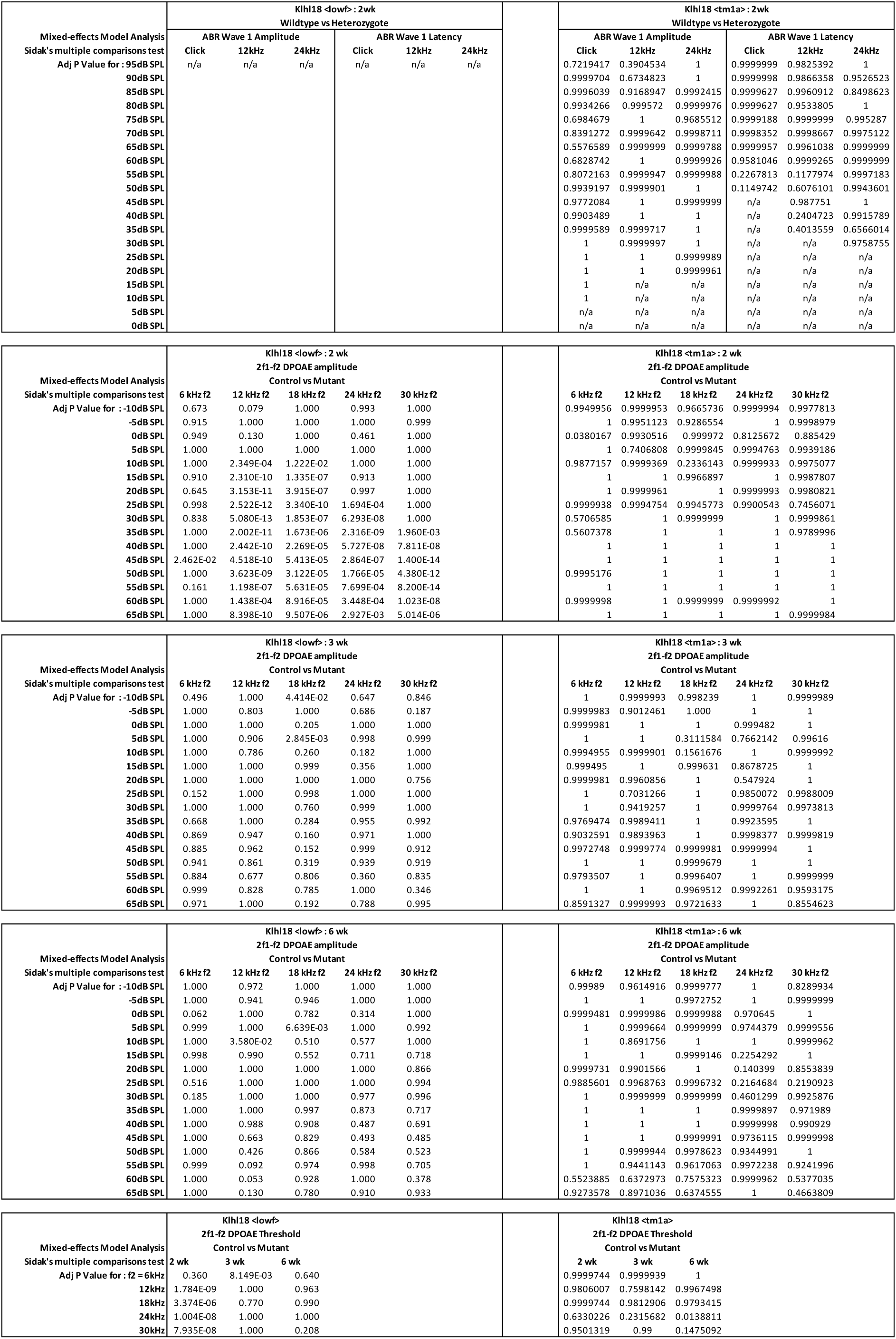
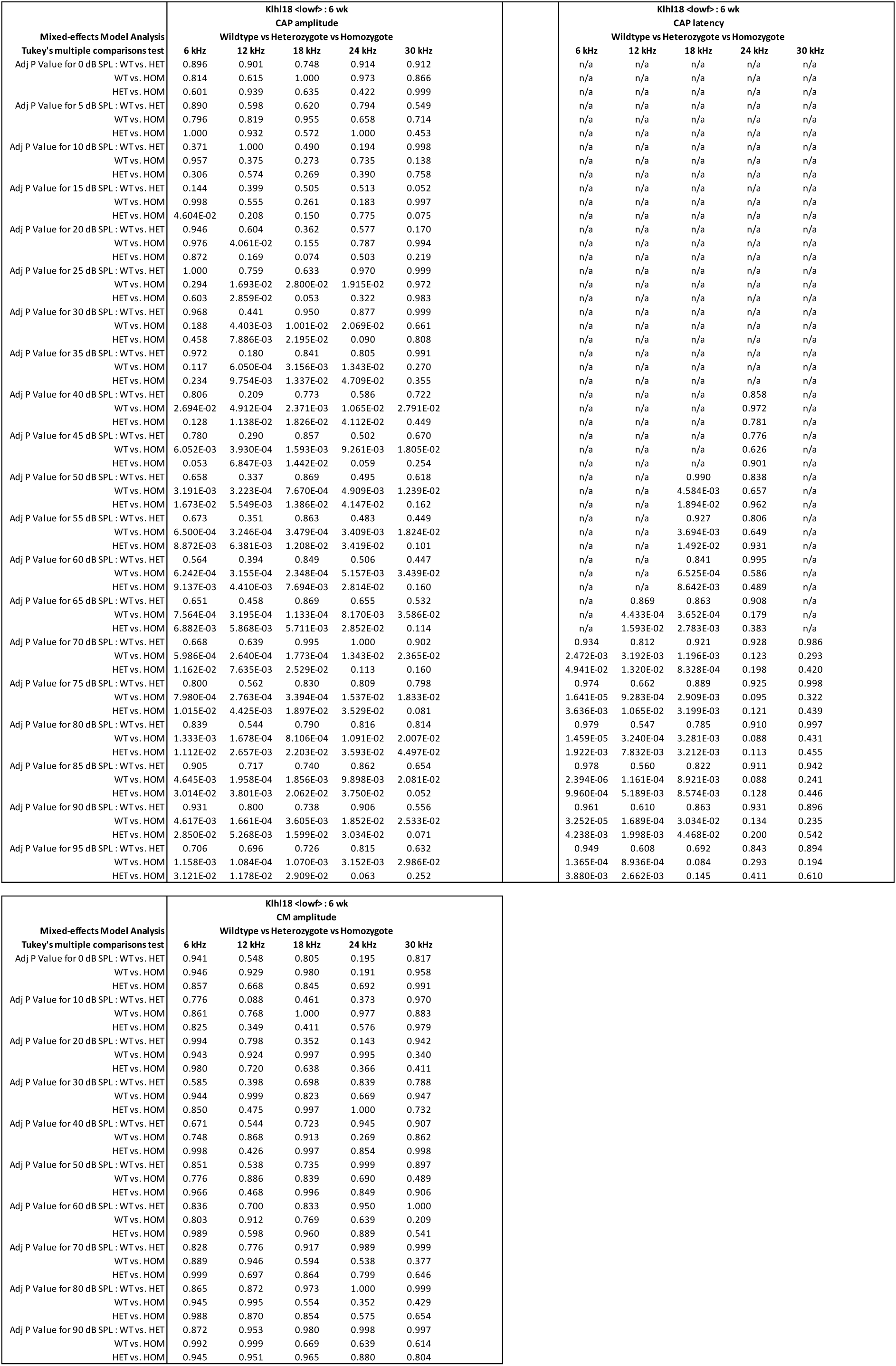
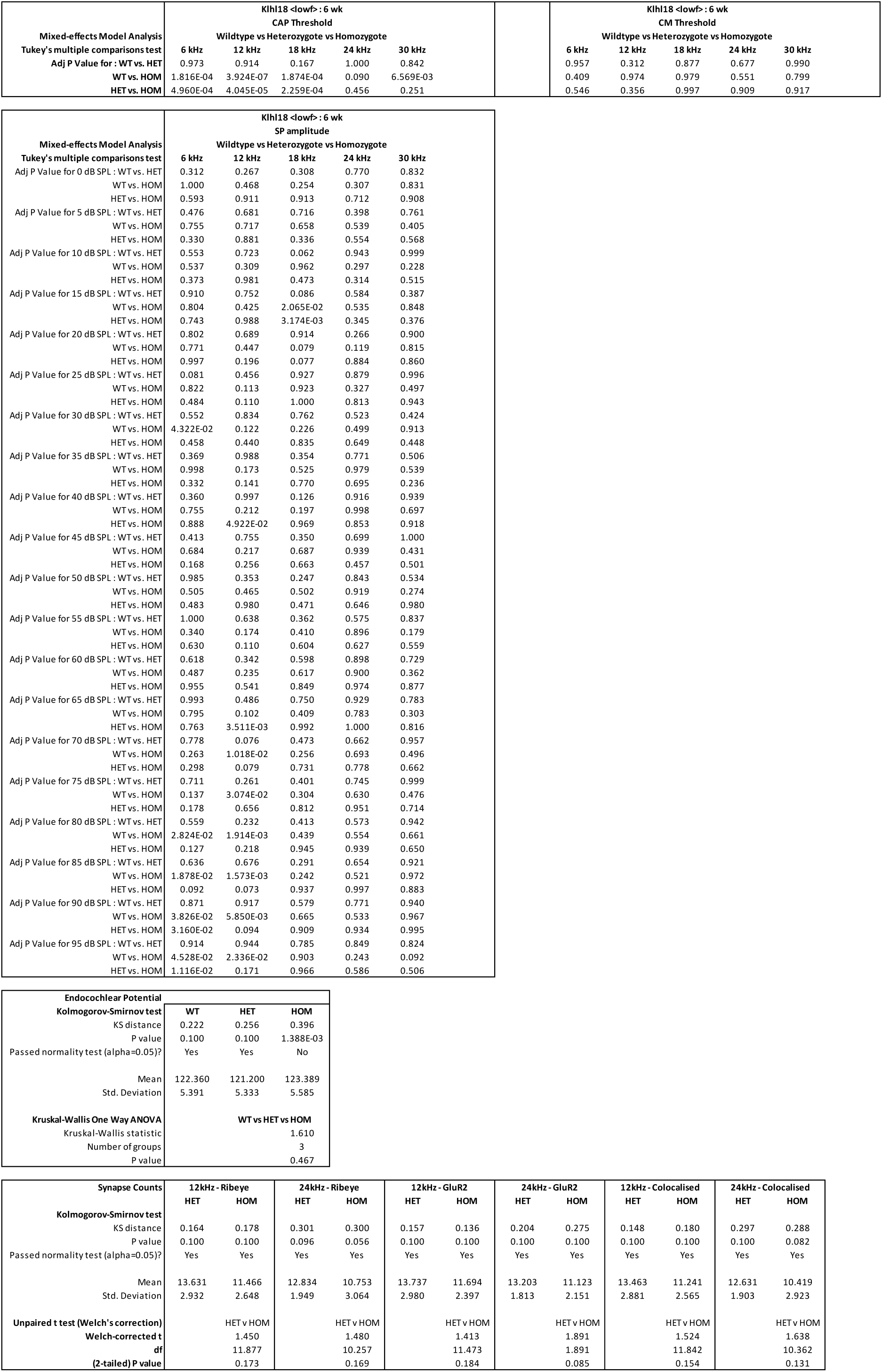
Additional Statistical Test Results.

## ABBREVIATIONS and UNITS

ABR: Auditory Brainstem Response
CAP: Cochlear Nerve Compound Action Potential
CM: Cochlear Microphonic
DAP: 4′,6-diamidino-2-phenylindole
dB: decibel
DPOAE: Distortion Product Otoacoustic Emission
EP: Endocochlear Potential
2f1-f2: the frequency of the DPOAE measured
f1: frequency 1, the lower stimulus frequency used to evoke DPOAEs
f2: frequency 2, the higher stimulus frequency used to evoke DPOAEs = 1.2 x f1
Fc: corner frequency (of a filter, digital or analog)
Hz: Hertz (cycles per second)
IHC: Inner Hair Cell
IOF: Input-Output Function
kHz: kiloHertz (x103 cycles per second)
µV: microVolt (x10-6 Volt)
mV: milliVolt (x10-3 Volt)
ms: millisecond (x10-3 seconds)
OHC: Outer Hair Cell
PBS: Phosphate-Buffered Saline
RWR: Round Window Response
SD: standard deviation of the mean
SEM: Scanning Electron Microscopy
SP: Summating Potential
SL: Sensation Level
SPL: Sound Pressure Level

## REFERENCES

Adams J, Kelso R, Cooley L (2000) The kelch repeat superfamily of proteins: propellers of cell function. Trends in Cell Biol. 10:17–24.

Albagli O, Dhordain P, Deweindt C, Lecocq G, Leprince D (1995) The BTB/POZ Domain: A new protein- protein interaction motif common to DNA- and actin-binding proteins. Cell Growth and Differentiation 6: 1193–1198.

Bespalova IN, Van Camp G, Bom SJ, Brown, Cryns K, DeWan AT, Erson AE, Flothmann K, Kunst HP, Kurnool P, Sivakumaran TA, Cremers CW, Leal SM, Burmeister M, Lesperance MM (2001) Mutations in the Wolfram syndrome 1 gene (*WFS1*) are a common cause of low frequency sensorineural hearing loss. Hum Mol Genet 10(22):2501–8.

Bowl MR, Simon MM, Ingham NJ et al (2017) A large scale hearing loss screen reveals an extensive unexplored genetic landscape for auditory dysfunction. Nat. Comm. 8:886.

Brandt A, Striessnig J, Moser T (2003) CaV1.3 Channels Are Essential for Development and Presynaptic Activity of Cochlear Inner Hair Cells. J. Neurosci. 23: 10832–10840.

Chaya T, Tsutsumi R, Varner LR, Maeda Y, Yoshida S, Furukawa T (2019) Cul3-Klhl18 ubiquitin ligase modulates rod transducin translocation during light-dark adaptation. EMBO J. 38: e101409.

Cody AR, Russell IJ (1987) The responses of hair cells in the basal turn of the guinea-pig cochlea to tones. J. Physiol. 383: 551–569.

Cryns K, Thys S, Van Laer L, Oka Y, Pfister M, Van Nassauw L, Smith RJH, Timmermans JP, Van Camp G (2003) The WFS1 gene, responsible for low frequency sensorineural hearing loss and Wolfram syndrome, is expressed in a variety of inner ear cells. Histochem. Cell Biol. 119:247–256.

Dhanoa BS, Cogliati T, Satish AG, Bruford EA, Friedman JS (2013) Update on the kelch-like (KLHL) gene family. Human Genomics 7:13.

Furness DN, Johnson SL, Manor U, Rüttiger L, Tocchettie A, Offenhausere N, Olt J, Goodyear RJ, Vijayakumarg S, Daic Y, Hackney CM, Franz C, Di Fiore PP, Masetto S, Jones SM, Knipper M, Holley MC, Richardson GP, Kachar B, Marcotti W (2013) Progressive hearing loss and gradual deterioration of sensory hair bundles in the ears of mice lacking the actin-binding protein Eps8L2. PNAS 110: 13989–13903.

Furukawa M, He YJ, Borchers C, Xiong Y (2003) Targeting of protein ubiquitination by BTB – Cullin3 –Roc1 ubiquitin ligases. Nature Cell Biol. 5: 1001–1007.

Harvey D, Steel KP (1992) The development and interpretation of the summating potential response. Hear. Res. 61: 137–146.

**Hunter-Duvar** **IM** (1978) A technique for preparation of cochlear specimens for assessment with the scanning electron microscope. Acto Oto-Laryngol. 85:sup351 3-23.

**Ingham** **NJ** (2019) Evoked potential recordings of auditory brainstem activity in the mouse: An optimized method for the assessment of hearing function of mice. Bio-Protocol 9: e3447.

Ingham NJ, Carlisle F, Pearson SA, Lewis MA, Buniello A, Chen J, Isaacson RL, Pass J, White JK, Dawson SJ, Steel KP (2016) *S1PR2* variants associated with auditory function in humans and endocochlear potential decline in mouse. Sci. Reports 6: 28964.

Ingham NJ, Pearson S, Steel KP (2011) Using the Auditory Brainstem Response (ABR) to Determine Hearing Sensitivity in Mutant Mice. Curr. Protocols. Mouse Biol. 1: 279–287.

Ingham NJ, Pearson SA, Vancollie VE, Rook V, Lewis MA, Chen J, Buniello A, Martelletti E, Preite L, Lam CC, Weiss FD, Powis Z, Suwannarat P. Lelliott CJ, Dawson SJ, White JK, Steel KP (2019) Mouse screen reveals multiple new genes underlying mouse and human hearing loss. PLOS Biol. 17: e3000194.

Kang MI, Kobayashi A, Wakabayashi N, Kim SG, Yamamoto M (2004) Scaffolding of Keap1 to the actin cytoskeleton controls the function of Nrf2 as key regulator of cytoprotective phase 2 genes. PNAS 101:2046–2051.

Kolla L, Kelly MC, Mann ZF, Anaya-Rocha A, Ellis K, Lemons A, Palermo AT, So KS, Mays JC, Hertzano R, Driver EC, Kelly MW (2020) Characterization of the development of the mouse cochlear epithelium at the single cell level. Nat. Comm. 11: 2389.

Kujawa SG, Liberman MC (2009) Adding insult to injury: cochlear nerve degeneration after "temporary" noise-induced hearing loss. J. Neurosci. 29: 14077–85.

Lewis MA et al. (2021) Collateral damage: Identification and characterisation of off-target mutations causing deafness from a targeted knockout programme. In Preparation.

Lopez-Escamez JA, Carey J, Chung WH, Goebel JA, Magnusson M, Mandalà M, Newman-Toker DE, Strupp M, Suzuki M, Trabalzini F, Bisdorff A (2015) Diagnostic criteria for Menière’s disease. J. Vestib. Res. 25:1–7.

Lynch ED, Lee MK, Morrow JE, Welcsh PL, León PE, King MC (1997) Nonsyndromic deafness DFNA1 associated with mutation of a human homolog of the Drosophila gene diaphanous. Science 278: 1315–8.

Mansour SL, Twigg SRF, Freeland RM, Wall SA, Li C, Wilkie AOM (2009) Hearing loss in a mouse model of Meunke Syndrome. Hum. Molec. Gen. 18: 43–50.

Melnick A, Ahmad KF, Arai S, Polinger A, Ball H, Borden KL, Carlile GW, Prive GG, Licht JD (2000) In- depth mutational analysis of the promyelocytic leukemia zinc finger BTB/POZ domain reveals motifs and residues required for biological and transcriptional functions. Mol. Cell. Biology 20: 6550–6567.

Minor DL, Lin YF, Mobley BC, Avelar A, Jan YN, Jan LY, Berger JM (2000) The Polar T1 interface is linked to conformational changes that open the voltage-gated potassium channel. Cell 102: 657–670.

Moghe S, Jiang F, Miura Y, Cerney RL, Tsai MY, Furukawa M (2012) The CUL3-KLHL18 ligase regulates mitotic entry and ubiquitylates Aurora-A. Biology Open 1:82–91.

Moser T, Starr A (2016) Auditory neuropathy – neural and synaptic mechanisms. Nature Reviews Neurology 12: 135–149.

Muller M, von Hunerbein K, Hoidis S, Smolders JWT (2005) A physiological place-frequency map of the cochlea in the CBA/J mouse. Hear. Res. 202: 63–73.

Ohn TL, Rutherford MA, Jing Z, Jung S, Duque-Afonso CJ, Hoch G, Picher MM, Scharinger A, Strenzke N, Moser T (2016) Hair cells use active zones with different voltage dependence of Ca^2+^ influx to decompose sounds into complementary neural codes. PNAS 113: E4716–25.

Pappa AK, Hutson KA, Scott WC, Wilson JD, Fox KE, Masood MM, Giardina CK, Pulver SH, Grana GD, Askew C, Fitzpatrick DC (2019) Hair cell and neural contributions to the cochlear summating potential. J. Neurophysiol. 121: 2163–2180.

Perez-Torrado R, Yamada D, Defossez PA (2006) Born to bind: the BTB protein-protein interaction domain. BioEssays 28:1194–1202.

Prosser HM, Rzadzinska AK, Steel KP, Bradley (2008) Mosaic complementation demonstrates a regulatory role for myosin VIIa in actin dynamics of stereocilia. A.Mol Cell Biol. 28:1702–12.

Reijntjes DOJ, Lee JH, Park S, Schubert NMA, van Tuinen M, Vijayakumar S, Jones TA, Jones SM, Gratton MA, Xia XM, Yamoah EN, Pyott SJ. (2019) Sodium-activated potassium channels shape peripheral auditory function and activity of the primary auditory neurons in mice. Sci. Rep. 9:2573.

Rhodes CR, Hertzano R, Fuchs H, Bell RE, Hrabé de Angelis M, Steel KP, Avraham KB (2004) A Myo7a mutation cosegregates with stereocilia defects and low-frequency hearing impairment. Mamm. Genome 15: 686–97.

Russell IJ, Cody AR, Richardson GP (1986) The responses of inner and outer hair cells in the basal turn of the guinea-pig cochlea and in the mouse cochlea grown in vitro. Hear. Res. 22:199–216.

Russell IJ, Legan PK, Lukashkina VA, Lukashkin AN, Goodyear RJ, Richardson GP (2007) Sharpened cochlear tuning in a mouse with a genetically modified tectorial membrane. Nat. Neurosci. 10: 215–23.

Sekerkova G, Richter C-P, Bartles JR (2011) Roles of the Espin Actin-Bundling Proteins in the Morphogenesis and Stabilization of Hair Cell Stereocilia Revealed in CBA/CaJ Congenic Jerker Mice. PLOS Genetics 7: e1002032.

Self T, Mahony M, Fleming J, Walsh J, Brown SD, Steel KP (1998) Shaker-1 mutations reveal roles for myosin VIIA in both development and function of cochlear hair cells. Development 125:557–66.

Skarnes WC et al. (2011) A conditional knockout resource for the genome-wide study of mouse gene function. Nature 474, 337–43.

Steel KP, Barkway C (1989) Another role for melanocytes: their importance for normal stria vascularis development in the mammalian inner ear. Development 107: 453–463.

Sun Y, Cheng J, Lu Y, Li J, Lu Y, Jin Z, Dai P, Wang R, Yuan H (2011) Identification of two novel missense WFS1 mutations, H696Y and R703H, in patients with non-syndromic low-frequency sensorineural hearing loss. J. Genet. Genom. 38: 71–76.

Taylor R, Bullen A, Johnson SL, Grimm-Gunter E-M, Rivero F, Marcotti W, Forge A, Daudet N (2015) Absence of plastin 1 causes abnormal maintenance of hair cell stereocilia and a moderate form of hearing loss in mice. Hum. Molec. Genetics 24: 37–49.

Velichkova M, Guttman J, Warren C, Eng L, Kline K, Vogl AW, Hasson T (2002) A human homologue of Drosophila kelch associates with myosin-VIIa in specialized adhesion junctions. Cell Motil Cytoskeleton 51:147–64.

Wan G, Corfas G, Stone JS (2013) Inner ear supporting cells: rethinking the silent majority. Semin. Cell Dev. Biol. 24:448–59.

Wan G, Corfas G (2017) Transient auditory nerve demyelination as a new mechanism for hidden hearing loss. Nat. Commun. 2017 8: 14487.

White JK et al. (2013) Genome-wide generation and systematic phenotyping of knockout mice reveals new roles for many genes. Cell 154: 452–64.

Xu L, Wei Y, Rebout J, Vaglio P, Shin TH, Vidal M, Elledge SJ, Harper JW (2003) BTB proteins are substrate-specific adaptors in an SCF-like modular ubiquitin ligase containing CUL-3. Nature 425: 316–321.

